# An integrated microfluidic and fluorescence platform for probing in vivo neuropharmacology

**DOI:** 10.1101/2024.05.14.594203

**Authors:** Sean C Piantadosi, Min‐Kyu Lee, Mingzheng Wu, Huong Huynh, Raudel Avila, Carina Pizzano, Catalina A Zamorano, Yixin Wu, Rachael Xavier, Maria Stanslaski, Jiheon Kang, Sarah Thai, Youngdo Kim, Jinglan Zhang, Yonggang Huang, Yevgenia Kozorovitskiy, Cameron H Good, Anthony R Banks, John A Rogers, Michael R. Bruchas

**Author notes:** These authors contributed equally (Sean C Piantadosi, Min‐Kyu Lee, Mingzheng Wu, John A Rogers, Michael R. Bruchas).

## Abstract

Neurotechnologies and genetic tools for dissecting neural circuit functions have advanced rapidly over the past decade, although the development of complementary pharmacological methodologies has comparatively lagged. Understanding the precise pharmacological mechanisms of neuroactive compounds is critical for advancing basic neurobiology and neuropharmacology, as well as for developing more effective treatments for neurological and neuropsychiatric disorders. However, integrating modern tools for assessing neural activity in large-scale neural networks with spatially localized drug delivery remains a major challenge. Here, we present a dual microfluidic-photometry platform that enables simultaneous intracranial drug delivery with neural dynamics monitoring in the rodent brain. The integrated platform combines a wireless, battery-free, miniaturized fluidic microsystem with optical probes, allowing for spatially and temporally specific drug delivery while recording activity-dependent fluorescence using genetically encoded calcium indicators (GECIs), neurotransmitter sensors GRAB_NE_ and GRAB_DA_, and neuropeptide sensors. We demonstrate the performance this platform for investigating neuropharmacological mechanisms in vivo and characterize its efficacy in probing precise mechanistic actions of neuroactive compounds across several rapidly evolving neuroscience domains.

## 1 INTRODUCTION

Using light to record and manipulate neural activity has dramatically advanced our understanding of the cell types (Boyden et al. 2005, Deisseroth and Schnitzer 2013), circuits, neurotransmitter systems, and receptors that give rise to complex mammalian behavior and contribute to pathological disease mechanisms. A key technique enabling these advances is fiber photometry (Cui et al. 2014), which records the fluorescence of biosensors, classically genetically encoded calcium indicators (GECIs) serving as a proxy of neuronal activity (Chen et al. 2013), with a high degree of spatial and temporal specificity (Simpson et al. 2024). More recent advances in biosensor development have led to the generation of fluorescent sensors for fast neurotransmitters (Marvin et al. 2013 2018 2019), monoaminergic neuromodulators (Feng et al. 2019, Patriarchi et al. 2018 2020, Jing et al. 2018, Sun et al. 2020, Wan et al. 2021, Zhou et al. 2023, Kagiampaki et al. 2023), neuropeptides (Duffet et al. 2022, Ino et al. 2022, Rappleye et al. 2022, Dong et al. 2022b), and intracellular signaling molecules (Lutas et al. 2022, Wang et al. 2022) that can be detected via fiber photometry in awake and behaving rodents. Combined with the comparatively low cost of fiber photometry (Simpson et al. 2024), the spatiotemporal specificity and every-growing toolkit of sensors for biological signaling molecules make it a valuable approach for investigating drug action mechanisms in freely moving mice, which can be typified by complex interactions and pharmacokinetic non-linearities (Wagner 1973). To date, no reliable method exists for combined site-specific drug delivery while locally monitoring continuous fluorescence biosensor signals, which could then be further enhanced with cell type, pathway, and receptor level specificity through advanced genetic tools (Stuber and Mason 2013).

Historically, the standard method to achieve a site-specific micro-injection requires an intracranially implanted cannula (Boschi et al. 1981). However, due to its large size, the conventional cannula system does not support the integration of optical neural recording modalities without substantially increasing the damage, inflammation, and significant lesioning of the brain tissue (Zhang et al. 2019, Wu et al. 2022). Furthermore, cannulation requires tethering an animal to an external pumping system that further limits their range of motion. Thus, there is a dearth of translational tools for simultaneously recording neural activity and localized fluid delivery of neuroactive compound. As a result, the precise cell type and receptor mechanisms mediating the therapeutic or off-target actions (including neurochemical alterations, direct and indirect effects on neural activity, and engagement of downstream signaling cascades) of many drugs remain elusive. Recently, interest in combining different or multiple methodologies to achieve local drug delivery and neural activity recording has grown. To achieve this, multiple approaches have been attempted, including multifunctional fibers for injections of viral vectors and neural recording (Park et al. 2017), electrophoretic drug delivery and electrophysiology recording (Proctor et al. 2018), and multi-shank neural probe (Yoon et al. 2022). Despite these substantial advances, devices retained several suboptimal features, including bulky construction, complexity in system design, and, most relevantly, the lack of fluorescence recording ability.

We have recently developed wireless, battery-free, fully implantable fluidic microsystems for user-controlled real-time intracranial photopharmacology (Wu et al. 2022). Here, we further adapt this device format to establish a multifunctional platform in which we demonstrate improved capabilities of intracranial drug delivery coupled with fluorescent biosensor recording in a spatiotemporally defined manner during spontaneous behavioral recordings. Miniaturized form factors of a drug fluidic microsystem (<0.15 g, 10 mm×13 mm) and flexible fluidic channels enable the integration of fluorescence recording modality into the platform for use across the whole mouse brain. The resulting system maintains the same ultralow-power operation, minimal heat generation, and battery-free functionality as previous device versions (Wu et al. 2022), but also includes additional capabilities in fluorescence recording through an integrated optic fiber. This system is: 1) customizable, allowing for different types of optic fiber with soft microfluidic channels that support versatility in types of recording and positioning across the brain; 2) capable of various modalities in a single platform, including recording or imaging, optical stimulation at different wavelengths, and simultaneous drug delivery with multiple compounds delivered independently; 3) compatible with a typical off the shelf, widely distributed fiber-photometry systems for detecting without the requirement of additional setups; 4) scalable with the use of lost-cost commercial components. Here, we provide several experimental demonstrations of these photo-fluidic devices using pharmacology and fluorescence recording and establish their utility for neuropharmacology experiments.

## 2 RESULTS

### Designs of optical fiber-integrated fluidic microsystem for direct modulation and monitoring of neural dynamics

The technology introduced here combines a standard fiber photometry system with a wirelessly programmable, battery-free electronic/microfluidic module (weight: <0.15 g, size: 10 mm×13 mm) (SI Appendix, Extended Data Fig. 1b) to support simultaneous neuropharmacology and fluorescent signal recording in small animal models (Fig. 1). The components include an electronics module and two micropumps (a cylinder shape: height 1 mm, diameter 2.45 mm) that interface to corresponding reservoirs (a hemispherical dome shape, capacity: 1.5 μL). Each independently connects to separate microchannels (two channels, each with a cross-sectional area of 30 μm by 30 μm) present in a thin, narrow, and mechanically compliant microfluidic probe (width 320 μm, thickness 150 μm, polydimethylsiloxane (PDMS)), bonded along the length of an optical fiber. Fluidic outlets (with cross-sectional areas the same as those of the channels) are positioned at the ends of the probe to allow real-time fluorescence measurements of neural activity changes in response to neuroactive compounds delivered by the microfluidics in the targeted region of the brain (Fig. 1b and SI Appendix, Extended Data Fig. 2a-c). Wireless energy harvesting and power management for the electronics follow from magnetic inductive coupling at a frequency of 13.56 MHz, as described previously for other different but related types of wireless devices (Shin et al. 2017, Yang et al. 2021). Software designed for present purposes forms a graphical user interface (GUI) to allow real-time control of the fluid pumps by triggering electrochemical reactions upon commands generated by a microcontroller in the electronic part of the module. The body of the device easily mounts on the skull using standard dental cement typically used for rodent neuro-hardware methods. The system setup appears in the SI Appendix, Extended Data Fig. 1a. Inlets located on the sides of each reservoir allow for refilling to enable multiple delivery events. This platform can support a wide range of experimental protocols, including the wide range of photometry applications described here but also pharmacology with multiple drug delivery epochs, fluorescence recording, optogenetics, and photopharmacology, with different types and lengths of optical fibers (SI Appendix, Extended Data Fig. 2d).

**FIGURE 1.**
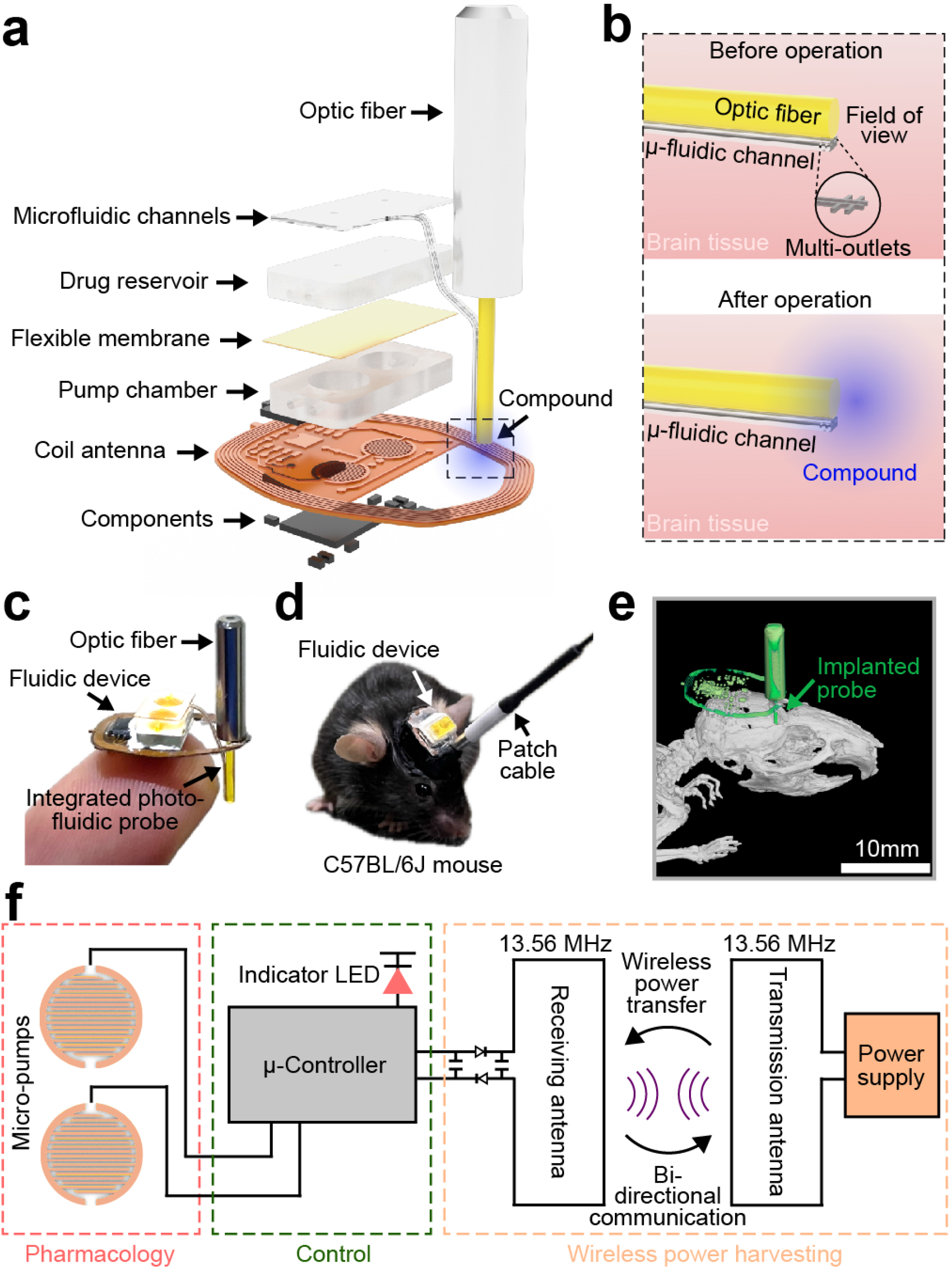
An integrated photo-fluidic microsystem for simultaneous pharmacology and fluorescence recording. (a and b) Exploded view illustration of the layer and the operation. (c) Picture of the photo-fluidic device on the tip of a finger (d) and after implantation into a WT mouse. (e) Picture and micro-CT image of mice with the devices implanted in the brain. (f) Electrical schematic diagram of the wireless, battery-free fluidic microsystem for programmable pharmacology.

### Design of an electrochemical micropump and microfluidic channel for dynamic flow control

Details of the microfluidic system design appear elsewhere (Wu et al. 2022), including data on its low power (<1 mW) operation, minimal heat generation (<0.2 °C), and significant driving force. Figs. 1a and Fig. 2a and b show the overall layout and the operation mechanisms. The hardware utilizes a lightweight, flexible printed circuit board (Cu/PI/Cu, 18 μm/75 μm/18 μm) that includes a coil antenna for wireless power transfer and control by near-field communication (NFC) protocols, mounting locations and interconnects for electronic components, and gold-coated interdigitated electrodes (Au/Cu, 200 nm/18 μm) aligned with pump chambers filled with an aqueous electrolyte solution (potassium hydroxide (KOH) 50 mM). An Au-coated flexible membrane of polystyrene-block-polyisoprene-block-polystyrene (SIS) separates the pump chambers from the overlying drug reservoirs. Commands issued through the GUI initiate hydrolysis reactions in the KOH solution (2H_2_O (liquid)

**FIGURE 2.**
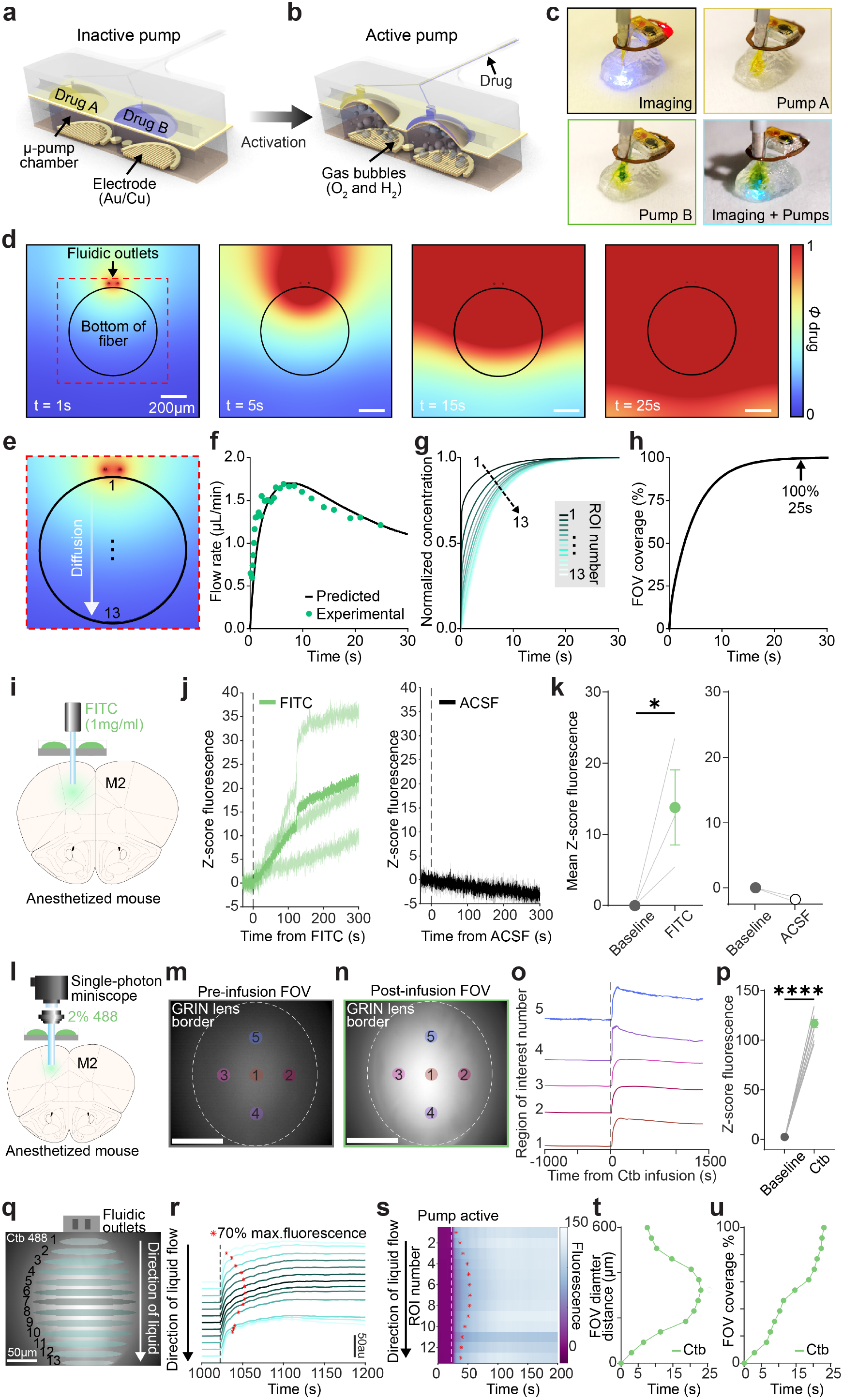
Operational features of opto-fluidic device, fluid dynamics modeling, and in vivo validation of optical detection of fluorescent dye delivery. (a) Schematic illustration of before and (b) after pump activation of the fluidic microsystem. (c) Various modes of operation. (d) Spatial distribution of the drug diffusion process in the brain at different times 1, 5, 15, and 25 sec showing the normalized concentration profile of the drug covering the optical probe. (e) High-magnification image of the probe of photo-fluidic device. 13 distinct regions along the diameter of the probe to quantify the timescale of FOV coverage. (f) Flow rate as a function of time (4 Hz, 50% duty cycle). (g) Normalized drug concentration as a function of time through different locations in the optical probe labeled from 1 – 13. (h) FOV Coverage (%) in the optical probe as a function of time. (i) Schematic illustration of the device with both drug reservoirs filled with FITC (1 mg/ml) dye. (j) Infusion of FITC (left) results in a rapid increase in fluorescence as measured by fiber photometry. Infusion of ACSF (right) does not change fluorescence. (k) FITC infusion significantly increases fluorescence relative to baseline (left; t(2)=3.55, p=0.04). ACSF infusion does not change fluorescence (right; p>0.05). (l) Schematic illustration of a GRIN lens-integrated device implantation into an anesthetized mouse with both drug reservoirs filled with fluorescent cholera toxin subunit B (Ctb) and miniature microscope. (m) FOV in M2 before pump activation. (n) Mean FOV after device operation and Ctb infusion. (Scale bar = 200 µm). Circles indicate manually added ROIs (1-5). (o) Fluorescence traces for ROIs identified in m and n. The dotted line indicates the activation of Ctb infusion. (p) Ctb infusion (green) results in a rapid and robust increase in fluorescence across the FOV (t(2)=27.1, p<0.0001). (q) Field of view from n with additional ROIs (1-13) spanning from the top of the FOV (closest to the fluidic outlets) to the bottom of the FOV (furthest from the fluidic outlets). Arrow indicates the direction of liquid flow. (r) Time-series fluorescence for each ROI in order from top of the FOV in q to the bottom (ROI 1-13) according to the distance from the fluidic outlet. ROIs are colored as in q. For visualization, time series are offset along the Y-axis by 25 arbitrary fluorescence units. Red asterisks indicate 70% of the maximal fluorescence value for each ROI time-series. (s) Time series data from r plotted as a heatmap, dotted line indicates pump activation. ROIs from q are ordered from top of the FOV to the bottom. Red asterisks indicate 70% of the maximal fluorescence value for each ROI time-series. (t) Plot of the time from the start of pump activation to reach 70% of maximal fluorescence for each ROI from the top (FOV diameter=0) to the bottom (FOV diameter=600) of the FOV. (u) FOV coverage (70% of maximal fluorescence) as a function of time from pump activation regardless of ROI location. (****p<0.0001, *p<0.05).

→O_2_ (gas) + H_2_ (gas)). Bubbles generated in this manner expand the volume of the pump chamber, thus mechanically deforming the SIS membrane. This deformation drives neuroactive compounds in the drug reservoir along the microfluidic channels of the probe to the brain. Multiple outlets associated with each microchannel allow the drug to spread evenly across the under the optical fiber (Fig. 2d). The Au coatings on the interdigitated Cu electrodes prevent degradation of the Cu due to electrochemical reactions with the KOH solution. The chemically inert and largely impermeable materials for the pumps and reservoirs allow for multiple cycles of use. The optical transparency of these materials allows visual access during drug-loading and device operation. A thin coating of SiO_2_ on the SIS membrane creates a strongly hydrophilic surface that avoids the formation of trapped air bubbles during the process of loading drugs into the system.

### Operation and fluidic characteristics of an integrated photo-fluidic device

Benchtop demonstrations of functionality involve a brain tissue model (a 0.6% agarose gel) that provides clear optical access to assess dynamic infusion by the device (Fig. 2c). The example illustrated here exploits fluorescence recording through an integrated optical fiber (Recording) before, during, and after a first (Pump A) and second (Pump B) activation, and fluorescence recording (Recording+Pumps). The operation of optical fiber with different wavelengths of light allows fluorescence recording of different biosensors and activation of opsins following pharmacological modulation. Finite element simulations of the optical probe and microfluidic channels show that the transient drug diffusion covers the bottom of the probe in approximately 25 seconds for an average drug delivery flow rate of 1.5 µL/min (Fig. 2d-h). Because recording fluorescence will occur in real time during behavior, obtaining a reasonable flow rate is necessary. Excessive rates can cause off-target effects and damage to brain tissue, thereby leading to low-quality recording through optical fiber. Inadequate rates not only increase the potential for blockage of the microfluidic channels but also lead to slow responses of limited interest for many experimental investigations (Zhang et al. 2019, Avila et al. 2021). An appropriate flow rate balances these two considerations to facilitate minimally invasive drug delivery evenly underneath the fiber. An adapted version of a previously reported system addresses requirements for present purposes (0.8 µL/min at 4 Hz and 10% duty cycle ≤ the appropriate range for the maximum flow rate ≤ 2 µL/min at 4 Hz and 50% duty cycle)(Wu et al. 2022). An intuitive GUI provides a real-time mechanism for controlling the rate through the frequency and duty cycle of the voltage applied to the electrolytic pumps (SI Appendix, Extended Data Fig. 3). For experiments here, these parameters are 4 Hz and 50% duty cycle, respectively, to ensure the maximum flow rates less than or equal to 2 µL/min for the robust but safe infusion. The analytical model shows a strong agreement with the experimental results (Fig. 2f) and predicts a temporal profile of delivery whereby the value of the maximum flowrate is 1.7 µL/min and occurs approximately 8 seconds after the drug delivery process initiates as the fluid travels from inside the drug reservoir, through the microchannels, and is ultimately delivered into the localized brain region. Diffusion of compound under the optical probe was determined at thirteen equally spaced points along the probe diameter (Fig. 2e, g, and h), showing the non-linear diffusion trends of the normalized water concentration reaching a steady state within 22-25 seconds. Similarly, the FOV coverage (%) was calculated based on the average normalized concentration in the area of the optical probe, taking 4.5 seconds to reach 63% coverage and 25 seconds to reach 100% coverage (Fig. 2h). Uniform drug delivery and diffusion towards the FOV enable chemical and optical modulation of neurons in the regions of interest, followed by high-quality recording of their spatiotemporal dynamics.

### Characterization of fluid infusion properties in vivo

Having now established successful operation of the photo-fluidic device ex vivo, we next evaluated whether spatially specific intracranial infusion could be reliably achieved with simultaneous fluorescence detection using a fiber photometry approach (Extended Data Fig. 5a). We anesthetized a mouse and implanted a photo-fluidic device (Extended Data Fig. 5b) into secondary motor cortex (M2) and loaded both drug reservoirs (1.5 µl) with fluorescein isothiocyanate (FITC; 1mg/ml), a green fluorescent dye (Fig. 2i), or artificial cerebrospinal fluid (ACSF). Following implantation, we operated the pumps of drug reservoirs at 10% duty cycle and 4Hz with anesthesia maintained. A dramatic change in fluorescence was detected immediately after activation of the pump containing FITC (Fig. 2j, left) compared to ACSF (Fig. 2k, right). Quantifying the mean fluorescence during the pre-infusion baseline and in the 5-minutes post-infusion, a significant increase in fluorescence was observed following FITC infusion (Fig. 2j, left) but not after control ACSF infusion (Fig. 2k, right). Infusion of the fluorescent retrograde tracer cholera toxin subunit B (CTB) produced a similarly robust increase in fluorescence (Extended Data Fig. 5d-f). These data indicate that our photo-fluidic device is, in principle, capable of producing rapid intracranial fluid delivery and that fluorescence underneath the delivery location is detectable by simultaneous fiber photometry.

Next, we quantified the spatial and temporal characteristics of the infused compound underneath the probe for fluorescence detection. We incorporated the flexible fluidic channels with a gradient refractive index (GRIN) lens, in combination with a single-photon miniature microscope, which produces images of the field of view underneath the probe during infusion. We again loaded both reservoirs with the green fluorescent compound CTB and implanted the GRIN fluidic device into M2 of an anesthetized mouse (Fig. 2l). Before the infusion, only tissue autofluorescence was detectable (Fig. 2m). However, following pump operation, the brightness of the FOV increased substantially (Fig. 2n; Supplementary Video 1) as CTB was infused. Fluorescence was rapidly and robustly increased across multiple regions of interest within the field of view (Fig. 2o-p). To better characterize the spread of liquid underneath the FOV in vivo, we examined ROIs across the FOV from closest to furthest from the fluidic outlets (Fig. 2q). We find that liquid rapidly travels across the FOV, initially covering the outer region of the FOV before spreading to the interior as quantified by the time needed to reach 70% of the maximal fluorescence of a given ROI (Fig. 2r-s). When plotted as a function of space underneath the fiber, this results in a gaussian spatial distribution of liquid across the FOV (Fig. 2t). The time course of the overall field of view coverage, regardless of ROI location across the FOV, was quite similar to our modeled data (Fig. 2g), with the full spatial occupation of the fluid cross the FOV achieved within 25s of pump activation (Fig. 2u). These data indicate that in vivo intracranial fluid delivery is robust, filling the entire area underneath the probe that detects fluorescence.

### Bidirectional manipulation of behavior and neural activity

With simultaneous fluid delivery and fluorescent detection capabilities established across several domains, we next determined whether the photo-fluidic device is capable of time-locked infusion of compounds with continuous fluorescence detection in vivo. We therefore sought to address whether the device could be used to monitor the effects of neural activity in response to drug delivery and then correlate these changes with behavioral effects in awake mice. We virally expressed the genetically encoded calcium sensor GCaMP6s in M2 neurons using pressure injection (Fig. 3a; Extended Data Fig. 6a). After four weeks of viral expression, we implanted the photo-fluidic device into M2 above our viral injection target before filling drug reservoirs with either ACSF, the α-amino-3-hydroxy-5-methyl-4-isoxazolepropionic acid (AMPA) receptor agonist AMPA (2mM in ACSF), or the GABA_A_-receptor agonist muscimol (0.75ng/nl) (Fig. 3b). Mice were then allowed to recover from anesthesia for 1 hour before being attached to a patch cable and being placed into a clear plexiglass chamber surrounded by a wireless RF antenna (Fig. 3c).

**FIGURE 3.**
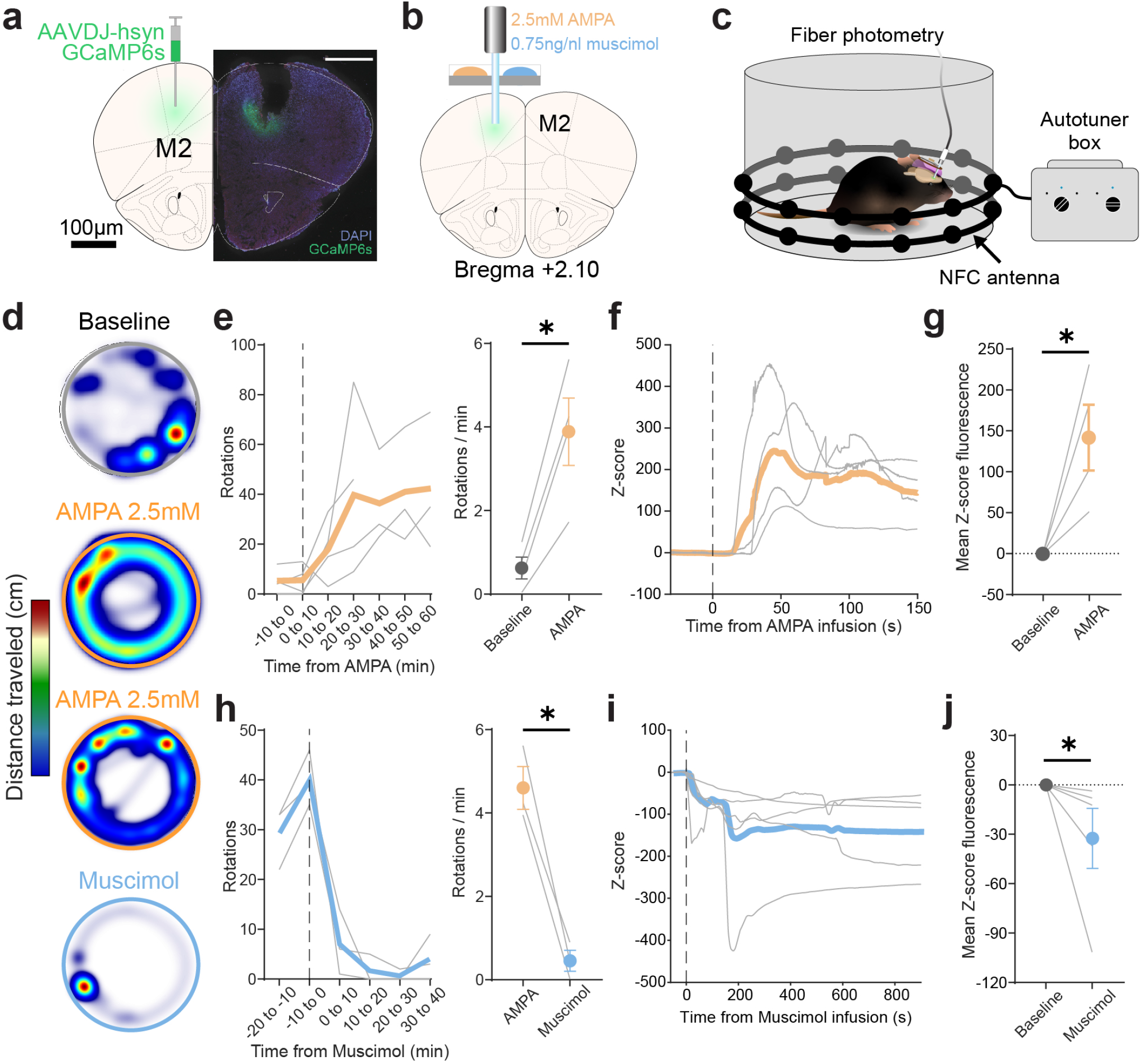
Bidirectional modulation of behavior and neural activity via photo-fluidic system. (a) Surgical schematic (left) and expression of GCaMP6s in M2 (right). (b) Experimental setup for α-amino-3-hydroxy-5-methyl-4-isoxazolepropionic acid (AMPA; orange) and muscimol (blue) infusions into M2. (c) Schematized image of photo-fluidic device operation and fiber photometry recording during behavior. (d) Locomotion heatmaps during baseline (grey outline), post-AMPA infusion (orange outline), and post-muscimol infusion (blue outline). (e; left) Number of rotations during 10 min time bins. The dotted line indicates AMPA (2.5mM) infusion into M2, leading to increased rotations. (e; right) AMPA infusion significantly increased the number of rotations per minute relative to baseline (t(3)=5.8, p=0.01). (f) Z-score fluorescence following AMPA infusion. (g) AMPA significantly increases mean fluorescence relative to baseline (t(3)=3.5, p=0.04). (h; left) Number of rotations during 10 min time bins. The dotted line indicates Muscimol (0.75 ng/nL) infusion into M2, reducing rotations. (h; right) Muscimol infusion significantly reduced the number of rotations per minute relative to AMPA infusion (t(2)=7.7, p=0.02). (i) Z-score fluorescence following Muscimol infusion. (J) Muscimol significantly reduces mean fluorescence relative to baseline (t(4)=2.3, p=0.044). *p<0.05.

Mice then underwent a 10-minute baseline during which they explored a circular arena while the spontaneous neuronal activity in the M2 was recorded (Fig. 3d). Subsequently, the pump containing AMPA was activated, resulting in unilateral drug infusion into M2. As expected following unilateral motor cortex activation (Wu et al. 2022), AMPA infusion produced a significant increase in rotational behavior (baseline mean rotations = 0.62 ±0.26SEM, AMPA mean rotations = 3.9 ±0.81SEM) where mice engaged in constant circling of the chamber that lasted for over an hour after infusion (Fig. 3e). Direct M2 infusion of AMPA produced dramatic and sustained increases in M2 GCaMP6 activity concomitant with the increased rotational behavior, as measured by the rise in fluorescence through the fiber optic (Fig. 3f; Supplementary Video 2). Relative to the pre-infusion baseline (baseline mean Z-score = -0.28, ±0.1SEM, AMPA mean Z-score = 141.7, ±40.17SEM), AMPA infusion produced dramatic increases in GCaMP6 fluoresence, indicating a large increase in M2 activity (Fig. 3g). By comparison, infusions of ACSF into M2 did not produce significant changes in locomotion, rotational behavior, or GCaMP fluorescence (Extended Data Fig. 6b-g), indicating that the observed increases in neural activity after AMPA infusion were due to agonist binding at ionotropic AMPA receptors resulting in excitatory engagement of M2 in a unilateral manner.

With the independent operational control of multiple drug reservoirs, we tested whether we could infuse a pharmacological agent that would reduce neural activity (muscimol) and examine whether the rotational behavior and hyperactivity induced by AMPA were re-normalized. In mice that received AMPA infusions, which resulted in robust rotational behavior as well as increased M2 activity, we then activated the second pump containing muscimol (Fig. 3b). Upon pump activation, the number of rotations immediately dropped to baseline levels (muscimol mean rotations = 0.46, ±0.25SEM) prior to AMPA infusion (Fig. 3e,h). Simultaneously, muscimol infusion resulted in a rapid and sustained reduction in M2 activity (Fig. 3i-j; Supplementary Video 3), indicating normalization of the hyperactive M2 activity. Together, these data establish that the photo-fluidic device generates bidirectional changes in neural activity and behavior in a time-locked manner following multiple types of drug infusion in a single animal.

### Combining site-specific pharmacology with optogenetics and neuromodulator sensing

The use of light to manipulate and measure neural activity has provided many insights into how neural circuits coordinate behavior (Simpson et al. 2024). More recently, the development of novel neurotransmitter biosensors have dramatically enhanced our understanding of their spatiotemporal dynamics. Multiplexing these neurotransmitter sensors with both light and drug delivery stands to provide answers to drug mechanisms of action and can be integrated with other optical approaches to reveal the endogenous function of neurotransmitter systems. To test whether this photo-fluidic system was capable of being combined with additional light delivery to manipulate activity during drug infusion, we focused first on a demonstration within the noradrenergic system, specifically the neuronal projection from the locus coeruleus (LC) to motor cortex (M2). Using mice expressing Cre-recombinase in neurons containing dopamine beta-hydroxylase (*Dbh*-cre), the enzyme responsible for converting dopamine into norepinephrine (NE), we first injected a virus encoding the red-shifted channelrhodopsin ChrimsonR (AAV5-DIO-ChrimsonR) into the LC followed by a virus encoding the green-shifted norepinephrine sensor GRAB_NE2m_ in M2 (Feng et al. 2019) (Fig. 4a, top). After four weeks of viral expression, a photo-fluidic device was implanted into M2 for of GRAB_NE2m_ fluorescence recording, optogenetic activation of LC terminals expressing ChrimsonR, and compound delivery (Fig. 4a, bottom; Extended Data Fig. 7a-b).

**FIGURE 4.**
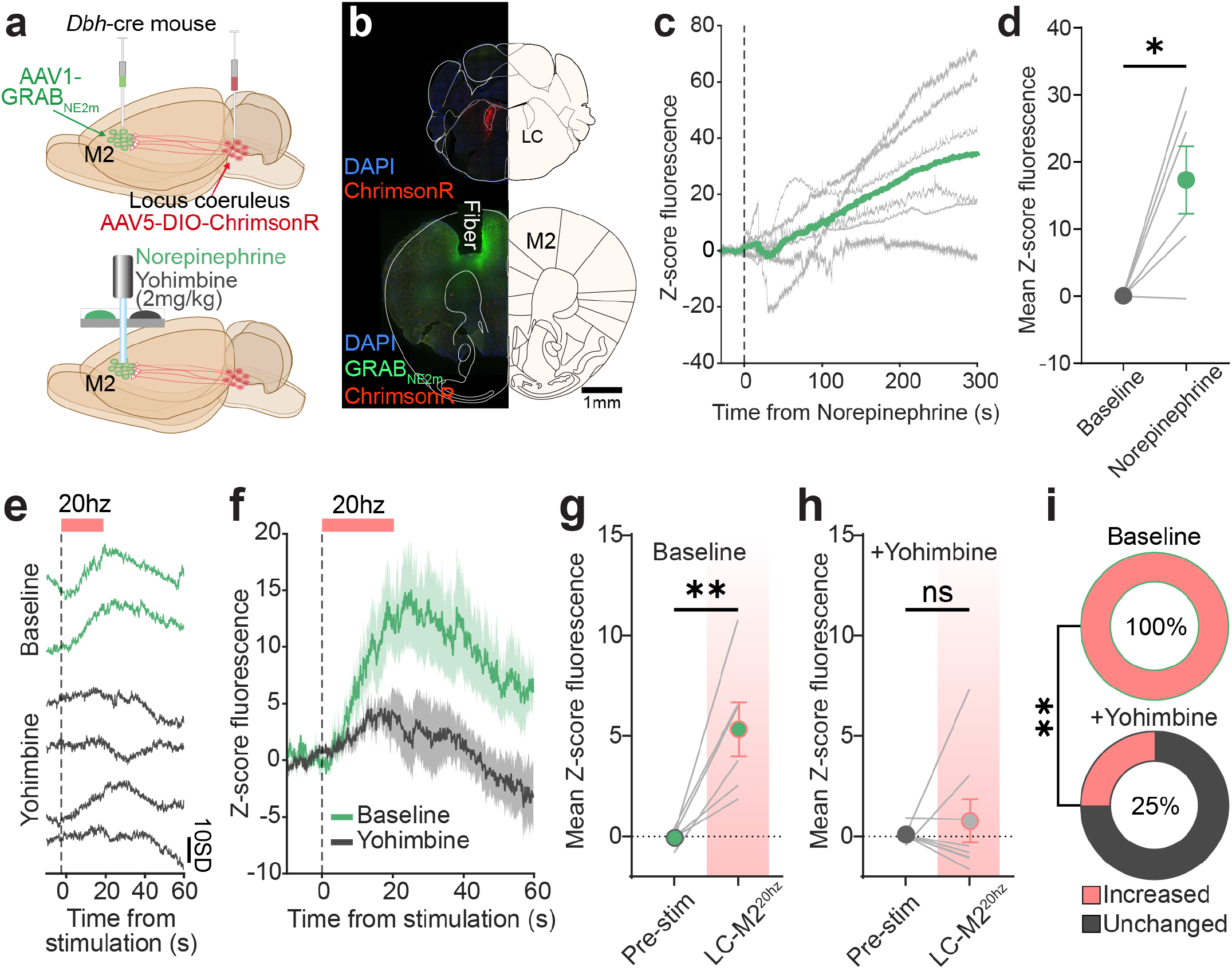
Multiplexing drug delivery with optogenetic activation locus coeruleus projections to M2. (a) Surgical strategy for optogenetic activation of locus coeruleus (LC) terminals in secondary motor area M2 with implant and infusion schematic below. (b) Histological verification of ChrimsonR expression in LC and GRAB_NE2m_ in frontal cortex below the optic fiber. (c) Infusion of norepinephrine (NE; 10-30mM) increases GRAB_NE2m_ fluorescence. (d) Infusion of NE increases GRAB_NE2m_ fluorescence relative to baseline (paired t-test, t(5)=3.45, p=0.02) (e) Representative individual trials from a single mouse during activation of LC-M2 (20hz, 20s) via an photo-fluidic device at baseline (green trace) and after infusion of the α2 antagonist yohimbine (0.3ng/1.5mL; grey trace). (f) Mean stimulation response at baseline and after yohimbine infusion of traces from e. (g) Mean Z-score fluorescence prior to infusion during baseline and during LC-M2^20hz^ stimulation across all trials (paired t-test, t(5)=4.13, p=0.009). (h) Mean Z-score fluorescence after yohimbine infusion during baseline and during LC-M2^20hz^ stimulation across all trials (paired t-test, t(7)=0.62, p=0.56). (i) Proportion of trials with increased (red) or unchanged (grey) GRAB_NE2m_ fluorescence following LC-M2^20hz^ stimulation at baseline (100% activated) and after yohimbine infusion (25% activated). Yohimbine significantly reduced the proportion of trials with increased GRAB_NE2m_ fluorescence relative to baseline (Chi-square test of proportions, X2=7.86, p=0.005).

We tested whether the infusion of norepinephrine (30µM in ACSF) into M2 would produce detectable changes in GRAB_NE2m_ fluorescence in M2. Infusion of NE into M2 resulted in sustained increases in GRAB_NE2m_ fluorescence (Fig. 4b; Supplementary Video 4). Compared to baseline GRAB_NE2m_ fluorescence, mean fluorescence following NE infusion was significantly elevated in all mice (Fig. 4c) Previously, this sort of calibration of receptor fluorescence in response to known concentration of ligand could only be conducted ex vivo. With this photo-fluidic device, the fluorescence response to known ligand concentration could then be directly compared with endogenous stimulus-evoked release or optogenetically evoked release to gain insight into the type or frequency of terminal activity that produces a given amount of neurotransmitter release in a specific brain region. To test whether this sort of optogenetically evoked response could be assessed and influenced pharmacologically using the photo-fluidic device, we conducted an experiment in which we optogenetically activated (20Hz for 20s) LC terminals expressing ChrimsonR in M2 as we recorded GRAB_NE2m_ fluorescence (Fig. 4a). Before any drug infusion, LC terminal stimulation at 20Hz elicited substantial increases in GRAB_NE2m_ fluorescence (Fig. 4d-e, green traces). We then activated the pump for a drug reservoir containing yohimbine (0.5µM in ACSF), an a2-adrenergic receptor (a^2^-AR) antagonist which will bind to GRAB_NE2m_, as it is a mutated a^2^-AR (Feng et al. 2019), and prevent NE from binding. Indeed, following infusion of yohimbine, 20hz stimulation no longer significantly increased GRAB_NE2m_ fluorescence in most trials due to receptor occupation by the antagonist (Fig. 4e-h). Consistent with these observations, 20hz terminal stimulation produced significant increases in GRAB_NE2m_ fluorescence on 100% of trials during baseline (Fig. 4i; top). By contrast, following yohimbine infusion only 25% of 20hz terminal stimulation trials resulted in increases in GRAB_NE2m_ fluorescence (Fig. 4i; bottom). These results demonstrate the compatibility of our photo-fluidic device for multiplexing sensitive pharmacology, projection-specific optogenetics, and neurotransmitter sensing in awake and behaving mice.

### Photo-fluidic measurement of interactions between neuromodulatory systems in the nucleus accumbens

Thus far, our investigations described here using the photo-fluidic device were limited to a relatively superficial brain region, M2. However, the majority of brain regions containing neurons that produce both monoaminergic neurotransmitters as well as neuropeptides are found in subcortical structures. Furthermore, we know that there are complex and poorly understood interactions between neurotransmitter signaling systems and neuropeptide signaling, often involving negative feedback loops (Ford 2014, Abraham et al. 2018). Traditionally, these interactions have been challenging to investigate in vivo at a systems level owing to a lack of tools to assess neuropeptide signaling with high spatiotemporal precision and the fact that most neuropeptides and related ligands do not readily cross the blood-brain barrier. Continued development of fluorescent sensors for neuropeptides has produced variants with high sensitivity and specificity for endogenous neuropeptides and exogenous ligands (Abraham et al. 2021, Wang et al. 2024). We propose that a device capable of precise spatiotemporal drug delivery and neuropeptide sensing would be able to provide significant insight into the complex interactions between peptide and neurotransmitter signaling systems in highly physiologically relevant and behavioral settings.

We next focused on the interaction between the κ-opioid receptor (KOR) system and dopamine (DA) signaling in the nucleus accumbens (NAc), a peptide signaling pathway known to regulate DA release (Tejeda and Bonci 2019, Ehrich et al. 2015, Escobar et al. 2020, Massaly et al. 2019). We first injected mice expressing Cre-recombinase in KOR-expressing neurons with a Cre-dependent, green fluorescent KOR-sensor kLight1.3b as well as a non-Cre dependent red fluorescent DA sensor, rGRAB_DA3m_ (Zhuo et al. 2023) (Fig. 5a; Extended Data Fig. 7c). After four weeks of viral expression, a photo-fluidic device was implanted into the NAc and drug reservoirs were loaded with DA (30µM) (Fig. 5a; Extended Data Fig. 7d). Mice were then attached to a patch cable, placed into a cylindrical plexiglass arena, and allowed to explore for at least 5 minutes. Green (470nm) fluorescence reflecting kLight1.3b activation and red fluorescence (560nm) reflecting rGRAB_DA3m_ activation were recorded simultaneously. After this baseline period, the pump containing DA was activated. DA infusion following pump activation produced locomotor hyperactivity, consistent with the effects of increased striatal DA (White et al. 2006, Woods and Ettenberg 2004)(Fig. 5b). Pump activation produced immediate robust and sustained increases in DA release as measured by rGRAB_DA3m_ fluorescence (Fig. 5c). Relative to the baseline period, DA infusion significantly elevated mean rGRAB_DA3m_ fluorescence across mice (Fig. 5d; Supplementary Video 5). We next examined whether elevating NAc DA levels with a DA infusion produced altered peptide binding to KOR as measured by kLight1.3b fluorescence. Interestingly, while DA infusion produced increases in rGRAB_DA3m_ fluorescence, a simultaneous reduction in kLight1.3b fluorescence was detected (Fig. 5e), such that across mice DA infusion increased mean rGRAB_DA3m_ fluorescence (Fig. 5f) and reduced kLight1.3b fluorescence (Fig. 5g). A correlation of rGRAB_DA3m_ and kLight1.3b fluorescence following DA infusion reveals that greater elevations in DA concentration corresponded to lower levels of kLight1.3b fluorescence, indicating that modest NAc DA concentration results in more dynorphin release relative to high DA concentration conditions (Fig. 5h). This same opposing pattern of rGRAB_DA3m_ and kLight1.3b fluorescence was observed when cocaine, which increases extracellular DA in the NAc (Stuber et al. 2005), was delivered systemically to these same mice (Extended data Fig. 7e-h). These data suggest interesting dynamic interactions between KOR and DA signaling in the NAc which have been previously suggested but not measured simultaneously (Ehrich et al. 2015, Shippenberg 2009). Visualizing the neuropharmacological effects of two receptor systems in parallel in an awake mouse could open many new avenues toward understanding fundamental interactions between neuromodulators and potentially identify novel therapeutic mechanisms.

**FIGURE 5.**
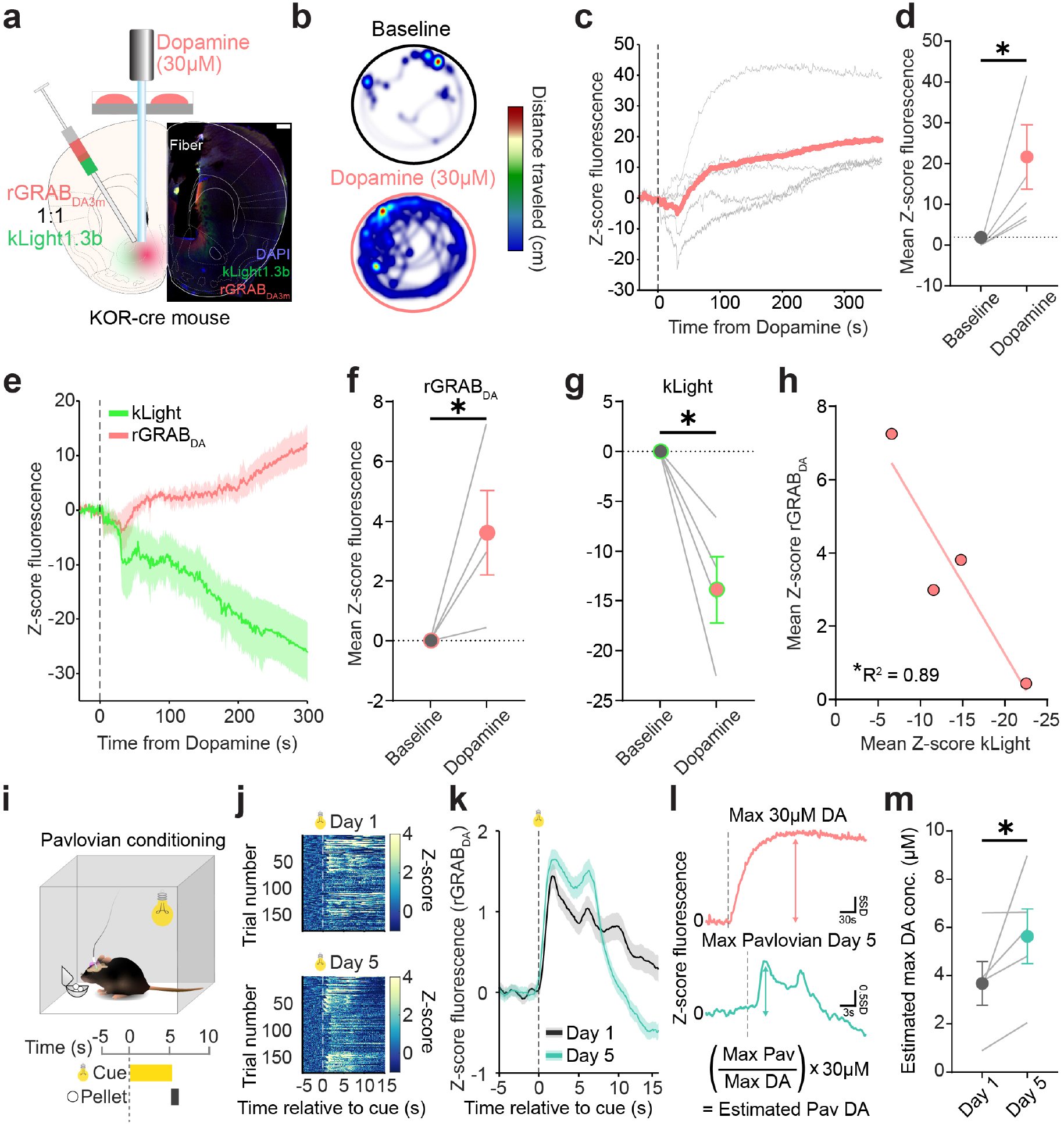
Photo-fluidic assessment of interactions between dopamine and dynorphin in the nucleus accumbens. (a) Schematic and histology depicting surgery co-injecting rGRAB_DA3m_ (red) and kLight1.3b (green) into the nucleus accumbens shell. Fluidic device for infusing dopamine (DA; 30 µM). (b) Representative heatmap of locomotion before (top) and after DA infusion (bottom). (c) Infusion of dopamine rapidly increases the fluorescence of rGRAB_DA3m_. (d) Infusion of DA significantly increases rGRAB_DA3m_ fluorescence relative to baseline (paired t-test, t(4)=2.5, p=0.03). (e) Infusion of DA produces an immediate increase in rGRAB_DA3m_ fluorescence and a simultaneous reduction in kLight1.3b fluorescence. (f) DA infusion into the NAc increases rGRAB_DA3m_ fluorescence (paired t-test, t(3)=2.6, p=0.04). (g) Significant reduction in kLight1.3b fluorescence following NAc DA infusion. (paired t-test, t(3)=4.16, p=0.013). (h) Significant negative correlation between rGRAB_DA3m_ fluorescence and kLight fluorescence following NAc DA infusion (Spearman correlation, r=0.93, p=0.04). (i) Schematic depicting pavlovian conditioning during fiber photometry several weeks after infusion. Mice received 40 trials of cue and pellet delivery per day across 5 days of training. A cue light lasting 5 seconds co-terminated with the presentation of a sucrose pellet. (j) Heatmaps of rGRAB_DA3m_ fluorescence aligned to cue onset for each trial during on Day 1 of training (top) and Day 5 of training (bottom). (k) Mean rGRAB_DA3m_ fluorescence traces aligned to cue onset across trials on Day 1 (grey) and Day 5 (cyan) of pavlovian conditioning. (l) Strategy for estimating concentration of stimulus-evoked DA release. Max Z-score fluorescence obtained following DA infusion and pavlovian conditioning. A ratio between these two values is obtained and then multiplied by the known concentration of DA infused to obtain an estimated max DA concentration evoked during pavlovian conditioning. (m) Effect of training on NAc DA concentration, with significantly elevated peak estimated DA concentration on Day 5 of pavlovian conditioning relative to Day 1 (paired t-test, t(4)=3.22, p=0.03). DA concentration was estimated for each individual animal using the maximum value observed during 30µM DA infusion in c. *p<0.05.

One potential additional utility provided by this photo-fluidic device is that a fluorescence response of a sensor to a known concentration of a ligand could be used to extrapolate and therefore infer ligand concentration that occurs in response to a stimulus or over the course of repeated behavioral events. Traditionally, estimates of monoamine concentrations using techniques such as fast-scan cyclic voltammetry (FSCV) were relative and subject to variability depending on the calibration technique used (Logman et al. 2000). By contrast, newly developed fluorescent sensors (with their known pharmacological constraints) could be combined with sensitive pharmacology of known ligand concentration, but exclusively in ex vivo preparations (Feng et al. 2019, Patriarchi et al. 2018, Sun et al. 2020). We believe our photo-fluidic device could be used to obtain relatively accurate concentration estimates in vivo. To investigate this possibility, we trained mice that previously received DA infusions on a pavlovian conditioning task where a conditioned stimulus (CS+; light cue) was paired with a sucrose pellet (US) over 5 days of training (Fig. 5i). During training, we recorded rGRAB_DA3m_ fluorescence and observed an evolution in the signal as a function of day (Fig. 5j-k). We then calculated a ratio between of the maximum value following infusion of 30µM DA (Fig. 5d) and the maximum value of the mean rGRAB_DA3m_ response across trials on Day 1 and Day 5 of pavlovian conditioning to obtain an estimate of the DA concentration evoked during pavlovian learning (Fig. 5l). Using these estimated values, we observed a significant increase in maximum DA concentration on Day 5 of training relative to Day 1 (Fig. 5m). Consistent with prior literature (Stuber et al. 2005), systemic administration of cocaine resulted in elevated estimated max concentration of relative to pavlovian conditioning days (Extended Data Fig. 7h). These data indicate that the concentration of spontaneously evoked transmitter release can be estimated in an intact animal and compared across behavioral paradigms subsequent to fluidic drug delivery.

## 3 DISCUSSION

In this article, we have demonstrated and characterized the utility of integrating intracranial drug infusion with optical recording of fluorescence, an increasingly popular and flexible modality for recording neurochemical and neurophysiological changes in vivo(Simpson et al. 2024), across a variety of potential use-cases. Construction of our photo-fluidic device is simple, inexpensive, and flexible, allowing for use with a variety of fiber-photometry systems (Fig. 1). We find that infusions are controllable, rapid, and robust, diffusing and covering the entire area under an optical probe (optic fiber or GRIN lens) within seconds of compound infusion (Fig. 2). Multiple drug reservoirs allow for spatiotemporally controlled bidirectional manipulation of neural activity as measured by calcium indicator fluorescence with simultaneous monitoring of behavioral response (Fig. 3). Our system also scales with the capabilities of the users’ photometric system and the exact experimental application. Here we demonstrate the capability of the photo-fluidic device to sense NE in vivo and it can be combined with optogenetic terminal activation and site-specific antagonist delivery (Fig. 4). Finally, we show that the photo-fluidic device can be implanted and operated deep within the brain to be used in conjunction with multi-colored fluorescent sensors for recording neuropeptide and neurotransmitter dynamics (Fig. 5). These proof-of-concept demonstrations provide a small glimpse of the biological insight that could be gained using the photo-fluidic device we present here.

This manuscript demonstrates several potential uses of the photo-fluidic device, though we note the flexibility of the device backbone allows for many additional applications utilizing intracranial fluid delivery. As with similar versions of these devices, injections of viruses can be achieved with the photo-fluidic device, ensuring accurate expression of viral expression underneath the optical probe (Jeong et al. 2015). A powerful potential use of this device could be to combine neuromodulator sensor photo-detection with cell-type specific calcium indicator expression, which has been done with numerous recording and imaging modalities (Zhuo et al. 2023, Lohani et al. 2022), with pharmacology to investigate the complex interactions between transmitter binding and changes in activity elicited by a drug of interest directly. This type of experiment bypasses complications that arise with systemic drug administration, including pharmacokinetic considerations, peripheral effects, and secondary effects interconnected brain regions (Howe et al. 2018). In addition, it allows for precise determination of ligand effects in a neural circuit of interest. Accurate assessment of neurotransmitter and neuropeptide concentrations elicited following endogenous release has traditionally been challenging in vivo. By using fluorescence values elicited with the infusion of known concentrations of ligand, estimates of concentration during spontaneous behavior can be made (Fig. 5m). These values can inform rates of drug clearance, receptor turnover, and other pharmacokinetic measures in an intact animal. Furthermore, pharmacological manipulation that alters these factors (e.g. infusion of an enzyme inhibitor) can be used to precisely quantify these complex interactions. Another pharmacological challenge that may be surmounted with this device is the accurate quantification of distinct monoamine release, which cannot be achieved with fast-scan cyclic voltammetry without the aid of pharmacology (Park et al. 2011 2009) or with new biosensors due to promiscuous binding affinities (Feng et al. 2019, Patriarchi et al. 2018, Sun et al. 2020, Zhuo et al. 2023, Kagiampaki et al. 2023). Here, site-specific infusion of known concentrations of a monoaminergic neurotransmitter (e.g. dopamine) could be infused while recording a norepinephrine biosensor to provide a quantitative assessment of the off-target binding. Likewise, this device also stands to scale with the rapid development of new biosensors (Dong et al. 2022a) and could be used to validate and compare sensors across generations during behavior. Finally, compounds that do not readily pass the blood-brain barrier can be directly infused with our photo-fluidic device, and their effects on neural activity and behavior can be evaluated in an awake and behaving mouse (Daneman and Prat 2015). Depending on the capability of a user’s fiber photometry system, the photo-fluidic device can also readily be used for photopharmacology (e.g. optopharmacology) experiments (McClain et al. 2023, Ahmed et al. 2023, Gómez-Santacana et al. 2022, Hüll et al. 2018), providing an additional level of control for more biological insights.

Recording and modulating neural activity in freely behaving small animals is critical for making inferences about how the brain controls behavior. An additional layer is the use of sensitive pharmacology to probe the receptors that influence the activity of a given neural population. This application can be very helpful for translational research directly related to treatment of neurological, neurodegenerative, and neuropsychiatric disorders. Despite its significance, there is a dearth of neuroscience tools capable of simultaneously monitoring neural phenomena (structural, neural activity, biochemical, and signaling molecule fluctuations) while locally modulating receptors pharmacologically. Previous research endeavors have addressed this challenge by integrating electrophysiological recording with drug delivery systems, enabling simultaneous recording and chemical modulation of neural activity. For instance, multifunctional neural probes have been developed, incorporating materials such as tungsten (Garwood et al. 2023, Ramadi et al. 2018), poly(3,4-ethylenedioxythiophene):PSS(PEDOT:PSS) (Proctor et al. 2018), and platinum electrodes into microfluidic systems (Shin et al. 2017). These experiments provide important insight into how drugs alter activity of groups of local neurons. However, these existing methodologies possess inherent limitations—they are capable of capturing a single effect of a drug, how it alters electrical activity, from groups of unidentified neuron populations near the electrodes. By combining fluidic drug delivery with fluorescence recording, we have expanded the possible outcomes that can be examined from neural activity to neurochemical and molecular signaling readouts, which can be multiplexed depending on a user’s fluorescence recording setup (Simpson et al. 2024). In addition, as compared to electrophysiological recordings, the entire suite of rodent genetics can be harnessed, with viruses expressed in discrete cell types, neural populations, and with projection-specificity.

While the photo-fluidic device has many noted advantages, several key limitations exist. First, device operation was limited to acute (≤ 1 day after device implantation) intracranial infusion due to clogging of the fluidic channels that occurred after one day of implantation, a known difficulty of biological implantation and challenging hurdle to overcome (Wu et al. 2022, Yoon et al. 2022). Despite this limitation, we believe our demonstrations barely scratch the surface of the information that could be gained even with relatively acute infusion time-points and these data can be used to inform subsequent in vivo experimentation (Fig. 5i-m, Extended Data Fig. 7). Ongoing experiments seek to develop a solution to increase the chronicity of device operation in vivo. A second minor limitation is that while drug delivery is wireless, preventing the complication of tethering to inflexible tubing, mice are still tethered via a patch cable attached to a commutator. Future experiments will integrate the photo-fluidic system with wireless fiber photometry (Burton et al. 2020), which will allow for entirely untethered recordings ideal for home-cage investigation or social behavior. Finally, we emphasize that our estimates of DA concentration (Fig. 5l) are relative and only shown to demonstrate the potential for calibration based on fluorescence responses to infusion of known ligand concentrations, previously not possible in vivo in a mammal (Feng et al. 2019, Patriarchi et al. 2020 2018, Wan et al. 2021, Zhuo et al. 2023). Experiments calibrating with infusions of multiple ligand concentrations would generate spontaneous release estimates of increasing accuracy. Regardless, this type of in vivo calibration to obtain accurate concentration estimates during spontaneous behavior may improve over other methods that rely on ex vivo calibration before or after an experiment, which can substantially vary and differ from experiment to experiment (Logman et al. 2000). Despite these limitations, we believe the current photo-fluidic device can provide novel insights into neuropharmacological mechanisms and lead to improved drug development.

Future improvements to allow for single-cell resolution by combining flexible fluidic drug delivery with GRIN or microprism lenses, and incorporation with wireless photometric recording (Burton et al. 2020) would allow for even greater spatial precision and mechanistic insight into drug action within the brain and its link to behavioral alterations. Incorporating biochemical sensing into this multimodal system will enhance neuropharmacological understanding by concurrently capturing freely moving mouse behavior, with neurotransmitter release and neural circuit activity spanning more molecular-cellular dimensions.

## Abbreviations

M2: secondary motor cortex
NAc: nucleus accumbens

## 4 METHODS

### Fabrication of the Electronics

Fabrication of the flexible printed circuit board (fPCB) began with laser ablation of a sheet of Cu/PI/Cu (18 µm/75 µm /18 µm; Pyralux, DuPont Inc.), using an ultraviolet (UV) laser system (ProtoLaser U4, LPKF Inc.), to pattern both sides with circuit traces, bonding pads, coil antennas, and interdigitated electrodes. Electroless plating formed a coating of Au (thickness: 200 nm) on the interdigitated Cu electrodes to prevent oxidation from electrochemical reactions of the Cu with the KOH solution in the pump chamber. An In/Ag low-temperature soldering paste (Indium Corporations Inc.) bonded the various electronic components to the fPCB. Additional details on the electronic circuitry design and components can be found in previous publication (Wu et al. 2022). A thin epoxy coating (Loctite Marine Epoxy, Henkel Inc.) reinforced these bonds to prevent delamination. A uniform layer of parylene (14 μm; Specialty Coating Systems Inc.) and an overlayer of PDMS formed by dip-coating (thickness: 200 μm) encapsulated the entire circuit to avoid fluid penetration.

### Fabrication of the Microfluidic Probes

Photolithography and deep ion etching defined patterns of relief on a silicon (Si) wafer in the geometry and dimensions of the microfluidic structures. Spin-coating formed a layer of PDMS with a thickness of 100 μm (10:1 elastomer to curing agent; Sylgard 184, Dow Corning Inc.) on this wafer. Inserting the samples into a vacuum chamber removed trapped air in the polymer precursor. Heating at 110 °C for 3 min cured the precursor into a solid, elastomeric form. Exposure to UV light for 3 min on water-soluble tape (ASWT-1, AQASOL Inc.) and treating a channel layer on Si wafer with a corona discharge (Electro-Technic Products Inc.) for 20 s allowed robust bonding between them upon contact, thereby facilitating mechanical release of the channel layer from the Si wafer to a piece of water-soluble tape. Spin coating and thermal curing of PDMS onto another Si wafer formed a capping layer (thickness: 50 μm). Simultaneous corona treatment for 20 sec on both a channel layer on water-soluble tape and a capping layer on Si wafer, followed by bonding and heat treatment on a hot plate at 110 °C for 30 min, formed closed microfluidic channels. A UV laser system (ProtoLaser U4, LPKF Inc.) defined the geometry of the microfluidic probe.

### Preparation of Flexible Membranes for the Pumps

Applying a mold release spray to a Si wafer (Ease Release 200, Mann Release Technologies Inc.) prepared it for spin-coating with a SIS solution dissolved in toluene (1 g/ml) (Sigma-Alrich Inc.). Thermal curing at 80 ºC for 30 min and then peeling from the wafer completed the formation of a solid elastomeric membrane. Electron beam evaporation (AJA International Inc.) formed a multilayer of Ti/Au/Ti/SiO_2_ (5 nm/50 nm/5 nm/20 nm) on the membrane to render a largely impermeable, hydrophilic surface.

### Fabrication of the Pump Chambers and Drug Reservoirs

A milling machine (MDX540 CNC Mill, Roland Inc.) defined a cylinder and dome shape on a block of cyclic olefin copolymer (COC; thickness: 1 mm) to form the pump chambers and drug reservoirs, respectively, with side filling ports (diameter: 350 μm) on each chamber and reservoir.

### Assembly of the Device

A schematic illustration of the steps for assembly of the devices appears in the SI Appendix, Extended Data Fig. 4. Aligning the pump chambers to the interdigitated electrodes on the fPCB substrate prepared them for bonding with a commercial sealant (Marine Adhesive Sealant Fast Cure 5200, 3M Inc.). The same sealant adhered the Au-coated SIS membrane to the base side of the reservoirs. Drying overnight enabled the bonding of this assembly to the pump chambers. Superglue (Instant adhesive, Loctite Inc.) bonded the optical fiber (diameter: 0.6 mm, length: 6 mm, Doric Inc.) to the edge of fPCB. The inlets of microfluidic channels aligned to the outlets of the reservoirs, secured with a pressure-sensitive adhesive (EL-8932EE, Adhesive Research Inc.). After aligning the outlets of the microfluidic channels to the plane of the FOV of the optical fiber, the same superglue bonded the channel along the length of the optical fiber. A razor blade trimmed excess PDMS beyond the end of the fiber and finished the device assembly.

### Validation of Drug Delivery in Benchtop and Theoretical Modeling

An experimental flow rate analysis relied on MATLAB (MathWorks Inc.) to calculate the volume of droplets infused from the outlet of microfluidic channels. A digital camera captured the release of blue-dyed droplets from the device operation using an RF system (NeuroLux Inc.) against a white background. A MATLAB code looped through each frame of the video to crop the area of the droplet, enhance the contrast between the droplet and background, and convert the frame to greyscale. Background subtraction of these frames with a greyscale threshold completely isolated the droplet. The code also measured the pixel length of the droplet diameter and a reference to calculate the volume of the droplet at each frame. The results enabled the calculation of the average flow rate by dividing the total volume by the time elapsed.

An analytical model for electrochemical fluidic microsystems derived from singular perturbation methods in was used to predict the average flow rate over time (Avila et al. 2021 2022). In the model, a combination of three non-dimensional parameters is used to characterize the drug delivery process and predict the flowrate (*dV ∗* /*dt∗*) given by and initial environmental pressure *P*_0_*=0.10, initial volume *V*_0_*=0.52, and microfluidic resistance *M*\*=0.002. The expression for the flowrate is based on two terms that dominate the delivery at different timescales shown by a fast and slow variable 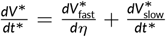. At the beginning of the delivery, the flowrate rapidly increases to reach the maximum value following the relationship:

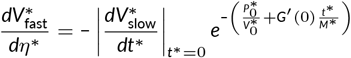

where 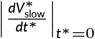 is a non-zero constant derived from the initial conditions. Then, after the maximum flowrate is reached, the flowrate transitions to a gradual decrease following. 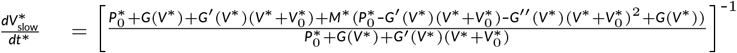 where the term *G*(*V*^*∗*^) is a non-dimensional pressure-volume function characterizing the mechanics of the flexible membrane and *G*^*′*^(*V*^*∗*^) and *G*^*′′*^(*V*^*∗*^) are the first and second derivatives of the function. Together, these two terms give the analytical relationship for the flow rate 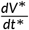.

Finite Element Analysis (FEA) was used to simulate the drug diffusion process in the brain and quantify the FOV coverage in the optical probe. A 2D diffusion model was developed in COMSOL, a commercial Multiphysics FEA software, to track the spatial concentration of the drug as it exits the microfluidic channels and evaluate the transient FOV coverage based on the range of drug delivery flow rates in the brain. The transient diffusion process was modeled as

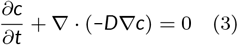

Where *D* is the diffusion coefficient (m^2^ sec^-1^), *c* is the drug concentration (mol m^-3^), and *t* is time (sec). The brain, optical probe (radius 0.6 mm), and square microfluidic channels (30 μm by 30 μm) were modeled using 115,000 triangular elements. A mesh convergence study was performed to ensure accuracy. Based on the optofluidic system’s operational conditions, an inward boundary flux of 1.5 µL/min was assigned to the microfluidic channel outlet to simulate the diffusion of water in the brain. The total simulation time was 50 seconds, with an output timestep Δ*t* of 0.2 seconds. The properties used in the simulation are *D*_tissue_ = 1 x 10^−9^ m^2^ sec^−1^, assuming a water concentration of *c*_0_ = 55 mol m^−3^. The 2D concentration contours capture the transient diffusion of the normalized concentration at the location of the optical probe and at 13 distinct regions along the diameter of the probe to quantify the timescale of FOV coverage (%) for drug concentrations greater than 70% in the range of the probe.

### Experimental Subjects

Adult male and female C57BL/6J, *Dbh*-Cre, and KOR-Cre mice were group-housed prior to surgery and given *ad libitum* access to food and water. Animals were maintained on a 12h:12h light:dark cycle (lights on at 9:00AM). All procedures were approved by the Animal Care and Use Committee of the University of Washington and Northwestern University and conformed to the US National Institutes of Health guidelines.

### Stereotaxic Surgery

For all surgeries, mice were anesthetized using 5% isoflurane mixed with oxygen and maintained at 1-2% isoflurane maintenance for the duration of the surgery. Mice were secured on a small-animal stereotactic instrument (Kopf Instruments, Tujunga, CA USA). Body temperature was maintained throughout surgery at 37°C using an infrared heating pad (Kent Scientific, Torrington, CT USA). Ophthalmic ointment was immediately applied to both eyes once mice were anaesthetized to prevent drying. Three rounds of betadine and alternating 70% ethanol wipes were applied to the scalp prior to an incision being made down the midline suture.

Surgical procedures were minorly different between experiments. For experiments involving fluorescent dye infusion into the frontal cortex (Fig. 2) entire experiment, a 1mm drill-bit was used to mice were under isoflurane anesthesia during the entire experiment, a 1mm drill-bit was used to create a craniotomy above M2 (coordinates A/P: +2.1, M/L: -1.1, D/V: -1.5 from bregma). A fiber- or GRIN-fluidic device was then lowered into M2. After reaching a depth of -1.5mm D/V, the skull was then dried and a layer of all purpose krazy-glue (Krazy Glue, Columbus, OH USA) was applied. After this application, a layer of black dental acrylic (Lang Dental, Wheeling, IL USA) was added to the skull and sides of the GRIN lens and built up into a secure base. After allowing 15 minutes for the dental cement to cure, pump reservoirs were filled with KOH (50mM) and drug reservoirs were loaded with either fluorescent cholera toxin subunit B, fluorescent FITC, or ACSF. The reservoir ports were then covered with kwik-seal and left to dry for 15 minutes. For fiber-fluidic experiments (Fig. 2i-k; Extended Data Fig. 5a-f) mice were then connected to a patch cable and fiber photometry system and pumps were operated to measure fluorescence changed induced by infusion. For GRIN-fluidic experiments (Fig. 2l-p), a miniature microscope (Inscopix nVista 3.0, Inscopix Palo Alto, CA USA) was lowered above the GRIN-fluidic device to allow for visualization of spatial diffusion of dye beneath the GRIN lens. The reservoirs were then prepared and operated as above. Mice for dye infusion experiments were transcardially perfused immediately after infusion.

For experiments involving drug infusion (Fig. 3, 4, & 5) mice underwent two separate surgeries. During the first surgery, mice received unilateral viral injections. All viruses were intracranially injected using a fixed-needle Hamilton Neuros syringe (Hamilton Company, Reno, NV USA) and an infusion pump (World Precision Instruments, Sarasota, FL USA) at a rate of 100 nl/min. In experiments recording activity of M2 neurons (Fig. 3), mice received a 500nl viral injection of GCaMP6f (AAV5-CAMKIIa-GCaMP6f-WPRE-SV40, Penn Viral Vector Core, Titer 1.31x10^13^) into M2 (coordinates A/P: +2.1, M/L: -1.1, D/V: -1.5 from bregma). For experiments recording norepinephrine sensor fluorescence in M2 while stimulating terminals from the LC (Fig. 4), mice received an injection of AAV1-Ef1a-GRAB_NE2m_ into M2 (coordinates A/P: +2.1, M/L: -1.1, D/V: -1.5 from bregma) and an injection of AAV5-hsyn-FLEX-ChrimsonR-tdTomato into the LC (coordinates A/P: -5.45, M/L: -1.1, D/V: -4.0 from bregma). For experiments recording dopamine and kappa opioid receptor sensor fluorescence in NAc (Fig. 5), mice received a combined 500nl viral injection (1:1 mixture) of AAV2/9-hsyn-rGRAB_DA3m_ (provided by Dr. Yulong Li) and pAAV-CAG-DIO-kLight into the NAc (coordinates A/P: +1.35, M/L: 0.65, D/V: -4.0 from bregma). Following these viral injections, skin was sutured using sterile silk sutures (McKesson Medical-Surgical, Irving TX, USA) and the viruses were left to express for at least 4 weeks.

Following viral expression, mice underwent a second surgery in which they were anesthetized as above and the previous craniotomy was identified and expanded if necessary. A fiber-fluidic device was then lowered into M2. After reaching a depth of -1.5mm D/V, the skull was then dried and a layer of all purpose krazy-glue (Krazy Glue, Columbus, OH USA) was applied. After this application, a layer of black dental acrylic (Lang Dental, Wheeling IL, USA) was added to the skull and sides of the GRIN lens and built up into a secure base. After allowing 15 minutes for the dental cement to cure, pump reservoirs were filled with KOH (50mM) and drug reservoirs were loaded and covered in kwik-seal. After kwik-seal was dry, mice were removed from anesthesia and allowed to recover for 1 hour prior to behavioral experimentation.

### Drugs

All drugs were dissolved in 1X artificial cerebrospinal fluid (ACSF; 31-33°C; 300-303 milliosmols) consisting of (in mM): 113 NaCl, 2.5 KCl, 1.2 MgSO4*g*.7H20, 2.5 CaCl2*g*.6H20, 1 NaH2PO4, 26 NaHCO3, 20 glucose, 3 Na+-pyruvate, 1 Na+-ascorbate. (RS)-AMPA hydrobromide (Cat 1074, PubChem ID 11957558) was purchased from Tocris Bioscience (Bio-Techne Corporation, Minneapolis MN, USA) and prepared at 2.5mM in 1X ACSF. Muscimol (Cat 0289, PubChem ID 4266) was purchased from Tocris Bioscience (Bio-Techne Corporation) and prepared in 1X ACSF at 0.75ng/500nl. Dopamine hydrochloride (Cat/Stock A11136) was purchased from Alfa Aesar (Ward Hill Massachusetts, USA) and prepared at 30µM in ACSF. L-(-)-Norepinephrine (+)-bitartate salt monohydrate (Norepinephrine; Cat A9512) was purchased from Sigma-Aldrich (Millipore Sigma, Burlington MA, USA) and prepared at 1mM and 10mM in ACSF. All drugs were made fresh and infused the same day they were prepared. Drugs were kept out of light to prevent oxidation and degradation.

### Behavioral apparatus and video recording

Animals were placed in a 30cm diameter cylindrical plexiglass arena with an antenna connected to a Neurolux radiofrequency system. During experiments recording GCaMP6 activity in M2 corncob bedding covered the bottom of the arena. For all other experiments no bedding was added to the plexiglass chamber. A Logitech webcam (Logitech, San Jose CA, USA) as well as a fiber-optic commutator were secured above the plexiglass arena to allow for free movement and behavioral recording. The TDT fiber photometry system was used to collect synchronous neural activity and video (30 FPS) recording.

### Device operation during behavior

Pump activation was manually operated by experimenters depending on the experiment using Neurolux software. Pump operation was conducted as previously described (Wu et al. 2022). A stable baseline period of at least 3 minutes was obtained prior to any pump activation for all experiments. Pump activation was manually timed-stamped live and confirmed via the top-down video recording. All infusions occurred < 6 hours after device implantation.

### Fiber photometry

Two LEDs were used to excite GCaMP6s. A 531-Hz sinusoidal LED light (Thorlabs, LED light: M470F3; LED driver: DC4104) was band-pass filtered (470 ± 20 nm, Doric, FMC4) to excite GCaMP6s and evoke Ca2+-dependent emission. For experiments recording red fluorescent rGRAB_DA3m_ (rGRAB_DA3m_) sensor activity in the NAc an additional 560nm LED was used. A 211-Hz sinusoidal LED light (Thorlabs, LED light: M405FP1; LED driver: DC4104) was bandpass filtered (405 ± 10 nm, Doric, FMC4) to excite GCaMP6s and evoke Ca2+-independent isosbestic control emission. Prior to behavior and recording, a 120 s period of GCaMP6s excitation with 405 nm and 470 nm light was used to remove most baseline drift. Laser intensity for the 470 nm and 405 nm wavelength bands were measured at the tip of the optic fiber and adjusted to *∼*70 μW before each day of recording. GCaMP6s fluorescence traveled through the same optic fiber before being bandpass filtered (525 ± 25 nm, Doric, FMC4), transduced by a femtowatt silicon photoreceiver (Newport, 2151) and recorded by a real-time processor (TDT, RZ5P). The envelopes of the 531-Hz and 211-Hz signals were extracted in real-time by the TDT program Synapse at a sampling rate of 1017.25 Hz.

### Optogenetic activation of LC terminals in M2

Optical stimulation of LC fibers expressing ChrimsonR in M2 was achieved through the same fiber used for recording GRAB_NE2m_ fluorescence. A 625nm LED was used for optogenetic stimulation, with the light power at the fiber tip set to 5mW. Optogenetic stimulation was manually triggered and delivered 20s long pulses of stimulation (20hz, 5ms pulse width). Mice received three separate 20s long stimulation trains during the baseline recording separated by three minutes. Three minutes after the final stimulation, yohimbine (0.3ng/1.5mL) was infused, followed by four more 20s stimulation pulses (20hz, 5ms pulse width).

### Pavlovian conditioning

One week prior to pavlovian conditioning experiments, mice were food restricted to 85% of their free feeding body weight and habituated to the reward (sucrose pellet) for 2 days prior to conditioning. Mice were habituated to handling, attachment to the fiber optic patch cable, and the experimental apparatus for 3 consecutive days before conditioning. Conditioning sessions took place in a conditioning chamber composed of transparent Plexiglas walls (30 cm × 30 cm × 30 cm) with a stainless steel grid floor (Med Associates, Fairfax, VT USA). A hole at the top of the plexiglass lid allowed for the patch cable connected to a commutator to be inserted into the conditioning chamber allowing for free exploration. A cue light (LED) was mounted on one wall of the chamber and served as the conditioned stimulus (CS). The unconditioned stimulus (US) consisted of a single 20 mg sucrose pellet delivered into a food hopper located on the opposite side of the chamber. Each conditioning session consisted of multiple trials, with a pseudorandomized inter-trial interval of 90 seconds. During each trial, the CS (cue light) was presented for 5 seconds, followed immediately by the delivery of the US (sucrose pellet). Mice underwent 5 days of conditioning with an average of 37 CS+US trials per day. rrGRAB_DA3m_ signal was recorded every other day of conditioning (Day 1, Day 3, Day 5).

### Fiber photometry data analysis

Custom MATLAB scripts were developed for analyzing fiber photometry data. The isosbestic 405 nm excitation control signal was subtracted from the 470 nm excitation signal to remove movement artifacts from intracellular Ca2+-dependent GCaMP6s fluorescence. Baseline drift was evident in the signal due to slow photobleaching artifacts, particularly during the first several minutes of each recording session. A double exponential curve was fit to the raw trace and subtracted to correct for baseline drift. After baseline correction, the photometry trace was z-scored relative to the mean and standard deviation of a baseline period prior to an event of interest. For pump activation, the baseline period consisted of the period prior to pump activation. For pavlovian conditioning, the baseline period was the period prior to CS+ onset. Quantification of peri-event fluorescence was calculated by taking the average of all samples during the baseline pre-event period (e.g. prior to pump activation) and comparing to the average of all samples during the post-event period (e.g. after pump activation).

### Estimates of DA concentration

To estimate DA concentration based off the response to infusion of known concentrations of DA (30µM), we took the maximum 560nm fluorescence response of rGRAB_DA3m_ and generated a ratio based off of the maximum mean response following the CS+ during pavlovian conditioning and the maximum response following systemic cocaine injection. This ratio was then multiplied by the DA concentration that was infused (30µM).

### Analysis of locomotion

Top-down videos of locomotion were fed into Noldus Ethovision (Noldus, Leesburg VA, USA) to analyze general locomotion (x/y location) as well as rotational behavior throughout each experimental session. A rotation was defined as 1 complete 360° turn clockwise or counter-clockwise with a 50° threshold.

### Histological verification of virus and probe placement

Mice were transcardially perfused with 10% formalin and phosphate buffered saline (1X PBS). Immediately after perfusion, heads (with implants intact) were placed into 10% formalin for 24h for post-fixation after which brains were removed and transferred to a 30% sucrose (in 1X PBS) solution. Brains were frozen and 35um sections were cut on a cryostat in 1:6 series. 1 series was mounted, counterstained with DAPI mounting media (Fluoroshield, Sigma-Aldrich) and imaged at 20x resolution on epifluorescence (Leica Microsystems, Wetzlar, Germany) to visualize virus and implant sites.

### Data analysis and statistics

Group statistical analyses were performed using GraphPad Prism 10 (GraphPad, LaJolla, CA). All mice were randomly assigned to experimental groups. Analyses were performed blind to experimental condition. For two-group comparisons, statistical significance was determined by either two-tailed independent samples t-tests or Mann-Whitney non-parametric tests depending on whether data were normally distributed.

For multiple group comparisons, repeated measures analysis of variance (ANOVA) was used followed by post-hoc analyses corrected for multiple comparisons. To compare proportions of trials with an evoked response, a Chi-square test of proportions was used. For all analyses, an α=0.05 was considered statistically significant.

## AUTHOR CONTRIBUTIONS

Conceptualization, S.C.P., M.K.L., M.W., J.A.R., and M.R.B.; Methodology, S.C.P., M.K.L., M.W., H.H., C.P., C.A.Z., Y.W., A.R.B., C.H.G., J.A.R., and M.R.B.; Theoretical simulation, R.A., and Y.H.; Investigation, S.C.P., M.K.L., M.W., H.H., C.P., C.A.Z., Y.W., J.K., S.T., and Y.K.; Writing – Original Draft, S.C.P., M.K.L., and M.W.; Writing – Review and Editing, S.C.P., M.K.L., M.W., R.A., C.P., C.A.Z., J.A.R., and M.R.B.; Funding Acquisition, Y.K., J.A.R., and M.R.B.; Resources, R.X., M.S., Y.H., Y.K., C.H.G., A.R.B., J.A.R., and M.R.B.; Supervision, Y.H., Y.K., C.H.G., A.R.B., J.A.R., and M.R.B.

## ACKNOWLEDGMENTS

This work made use of the NUFAB facility of Northwestern University’s NUANCE Center, which has received support from the SHyNE Resource (NSF ECCS-2025633), the IIN, and Northwestern’s MRSEC program (NSF DMR-2308691). We thank Dr. Azra Suko for lab management and organization, and Bailey Wells and Valerie Lau for assistance with the mouse colony. We also thank the entire Bruchas laboratory as well as the Neuroscience of Addiction, Pain, and Emotion (NAPE) Center at the University of Washington for resources alongside critical feedback.

## DATA AVAILABILITY

The data supporting this study’s findings are available upon request from the authors.

## SUPPORTING INFORMATION

Additional supporting information may be found in the online version of the article.

## SUPPLEMENTAL FIGURES

**Extended Data Figure 1:**
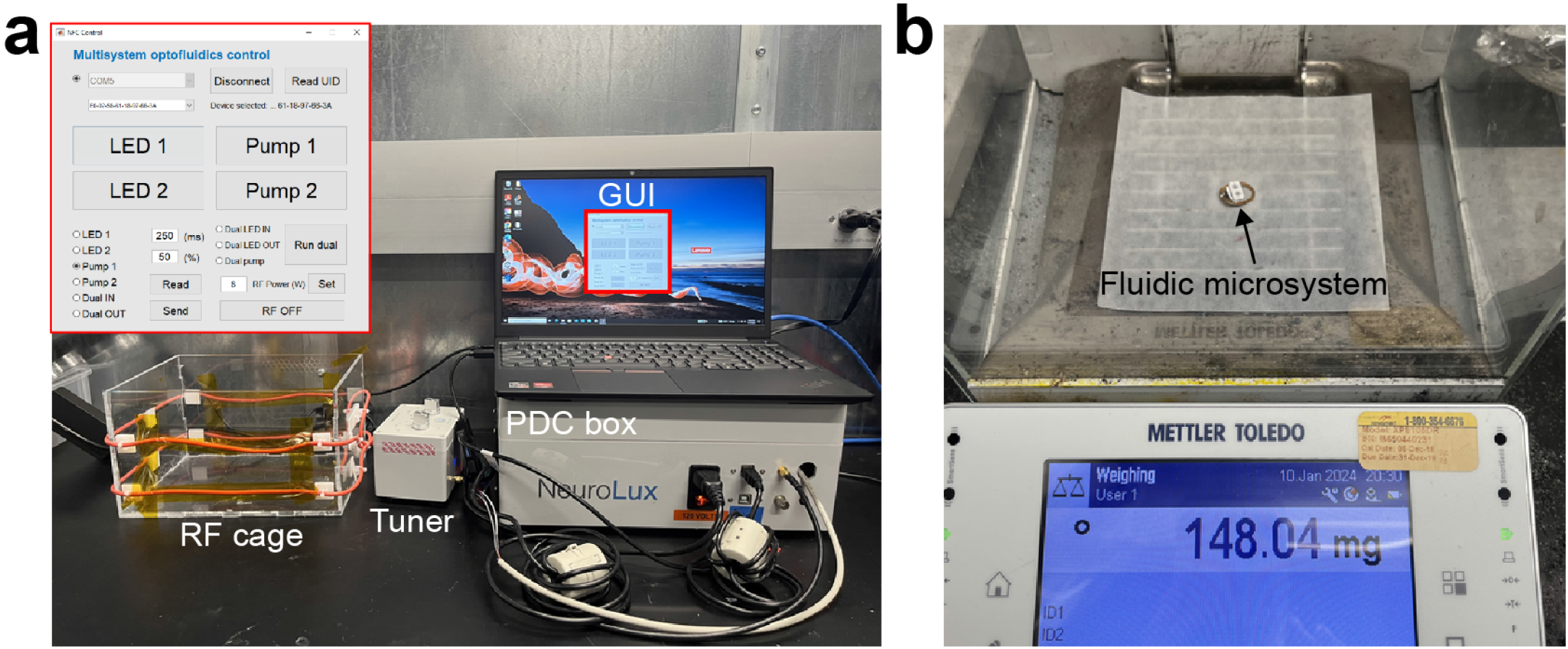
Images of the control electronics and the device. (a) Pictures of the wireless powering and communication system, including the RF cage, tuner, PDC box, and the graphical user interface. (b) Picture of a device on a scale to illustrate its weight.

**Extended Data Figure 2:**
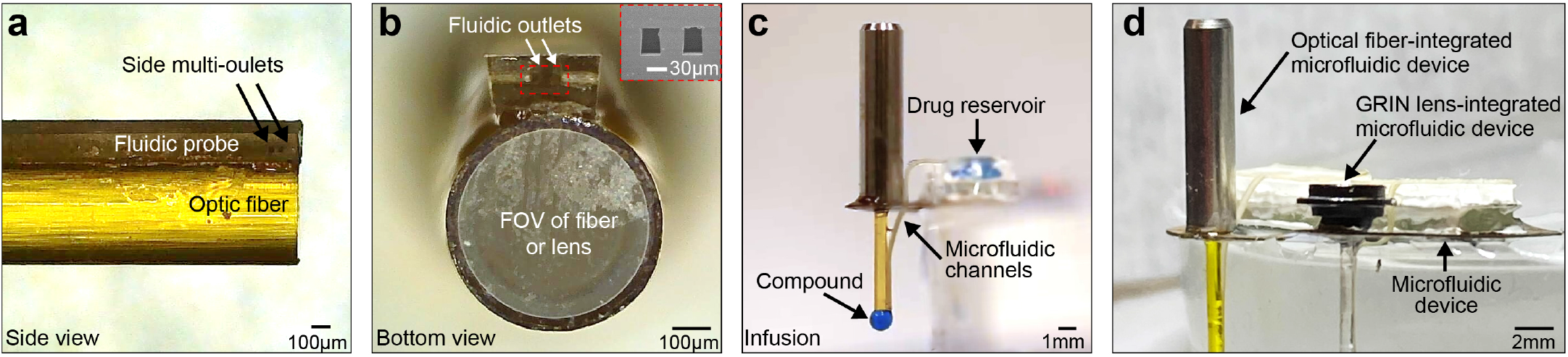
Pictures of optical fiber integrated with fluidic probe. (a) Structure of integrated photo-fluidic probe from the side view. (b) Photo of bottom of photo-fluidic device depicting bottom of fiber and fluid outlets. Inset represents scanning electron microscopy image of fluid outlets. (c) Operation of pumps and delivery of the compound to the under the optical fiber after device operation. (d) (left) Optical fiber-integrated microsystem for 1-photon fiber photometry and (right) GRIN lens-integrated microsystem for 2-photon imaging and miniature microscope.

**Extended Data Figure 3:**
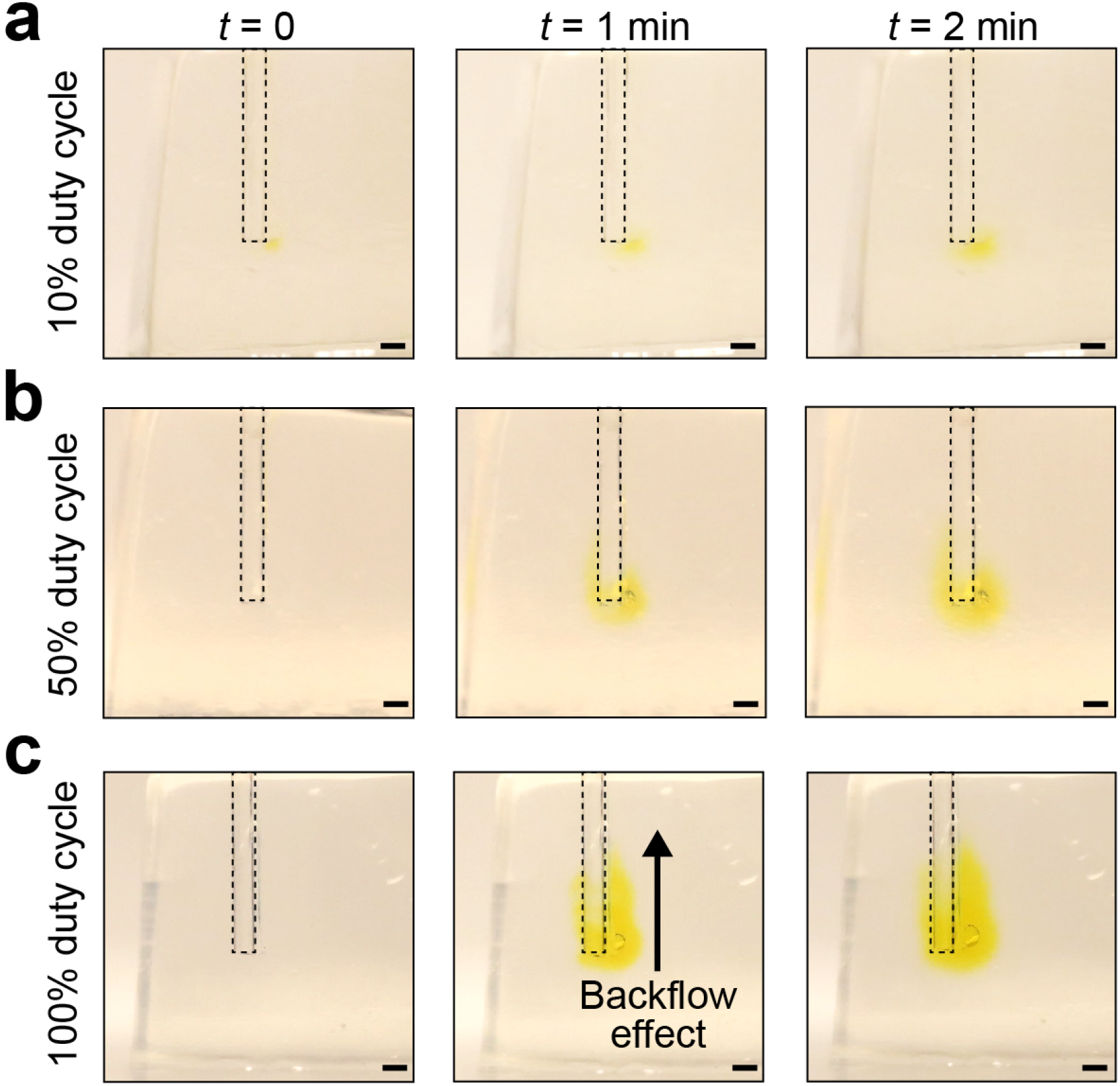
Pictures from an infusion test that uses a dyed (yellow) solution and a brain phantom (0.6% agarose hydrogel) with devices operated at different duty cycles. (a) 10%, (b) 50%, and (c) 100% duty cycle with 4 Hz frequency. (Scale bar = 1 mm). The dashed lines indicate the location of the optical fiber / microfluidic probe. The figures demonstrate that low-duty cycles (10% and 50%) with low flow rates can reduce the effects in which flow occurs along the interface between the probe and the surrounding phantom (black arrow).

**Extended Data Figure 4:**
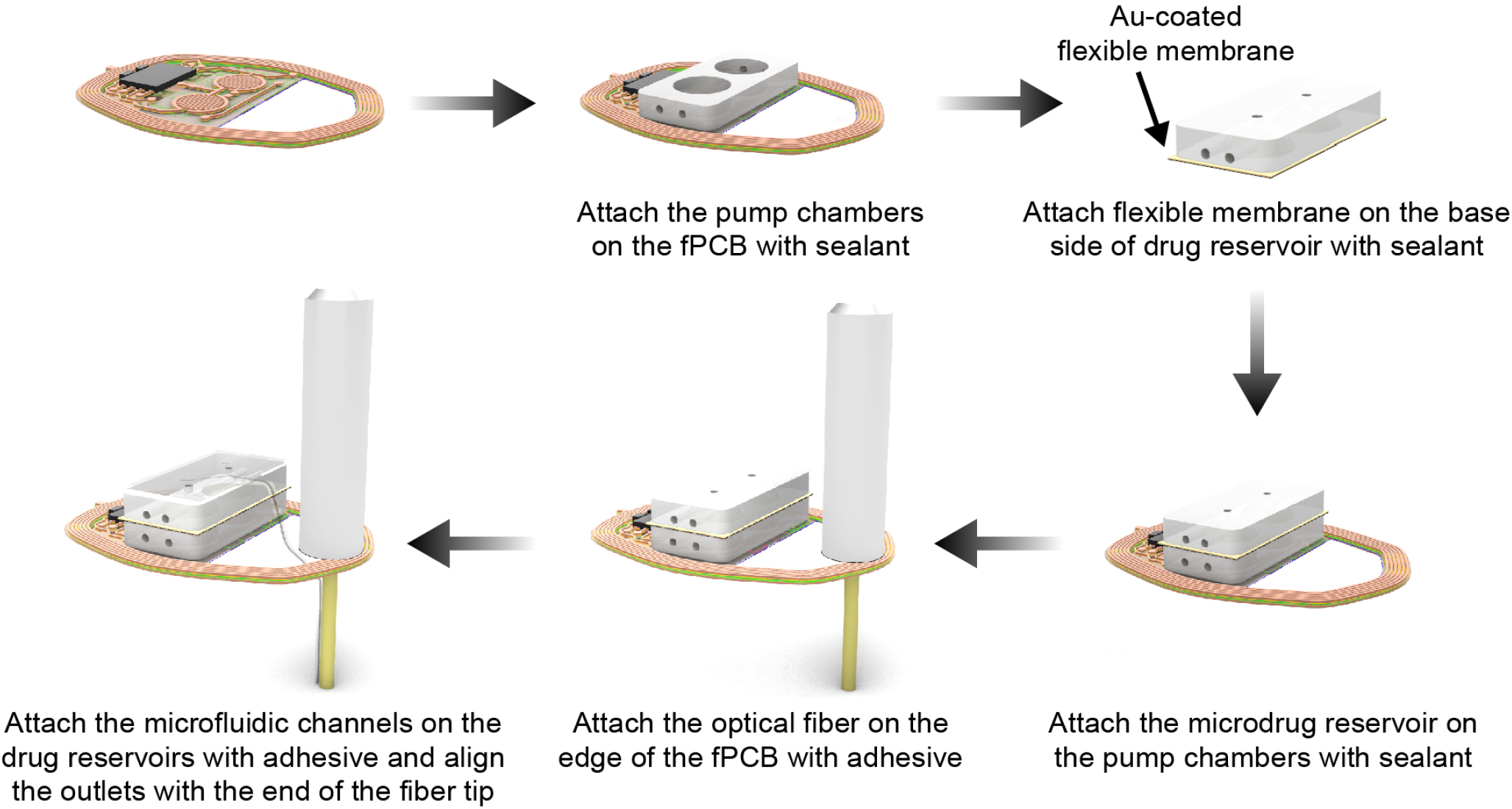
Schematic illustration of the steps for assembly of the devices.

**Extended Data Figure 5:**
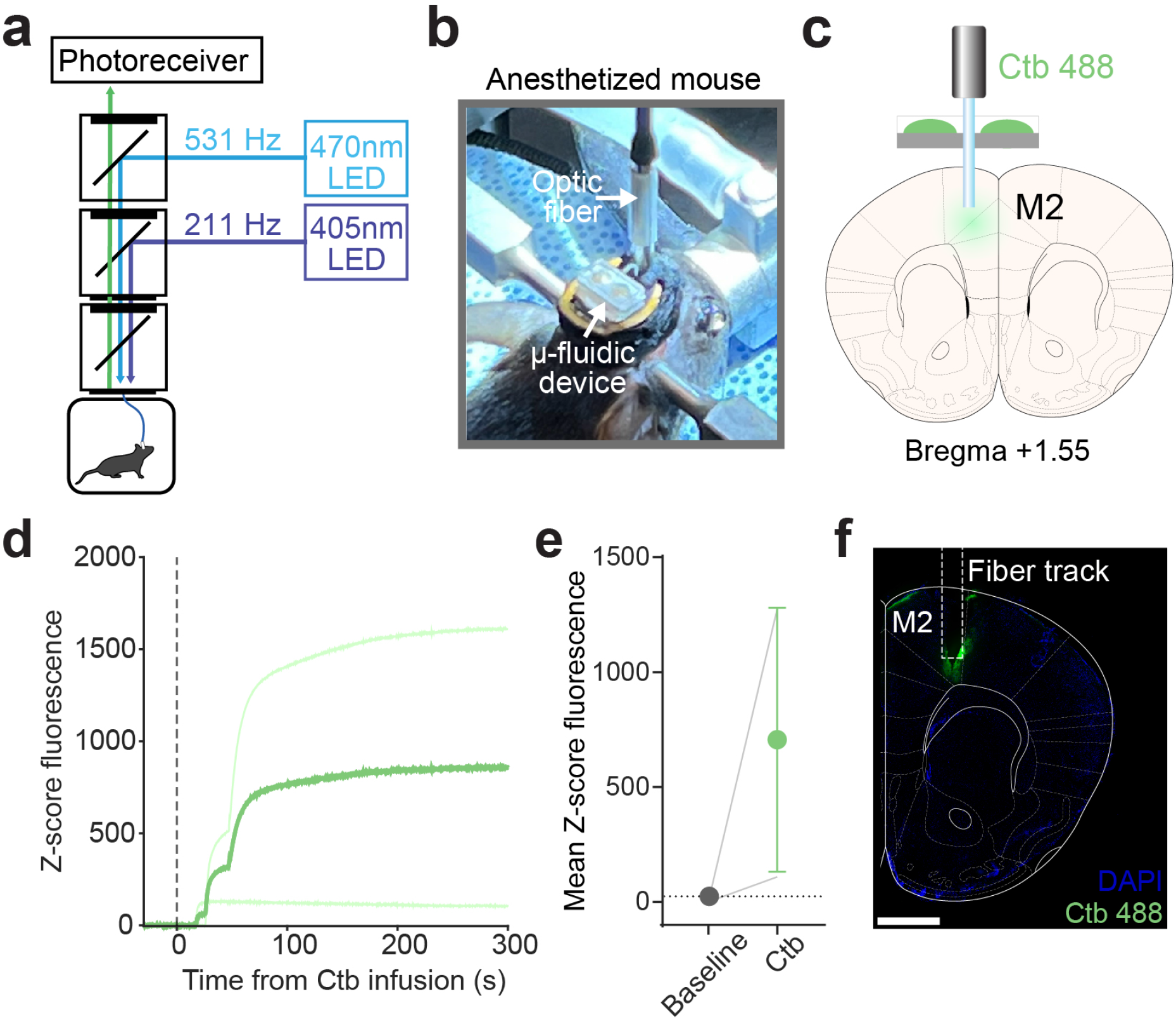
Infusion of Ctb into M2. (a) Depiction of fiber photometry system and LED paths. (b) Picture of the anesthetized mouse (isoflurane 1.5%) and fiber optic patch cable attached to the photo-fluidic device. (c) Schematic illustration depicting infusion of Ctb 488 into M2 of an anesthetized mouse. (d) Z-score fluorescence in response to Ctb 488 infusion. Dotted line indicates pump activation. Light green lines indicate individual infusions, dark green line denotes mean response across trials. (e) Mean Z-score fluorescence before and in the 5 minutes after Ctb 488 infusion. (f) Histological image showing Ctb 488 (green) expression in M2 following infusion. Photo-fluidic fiber track indicated by dotted rectangle. Scale bar = 1mm.

**Extended Data Figure 6:**
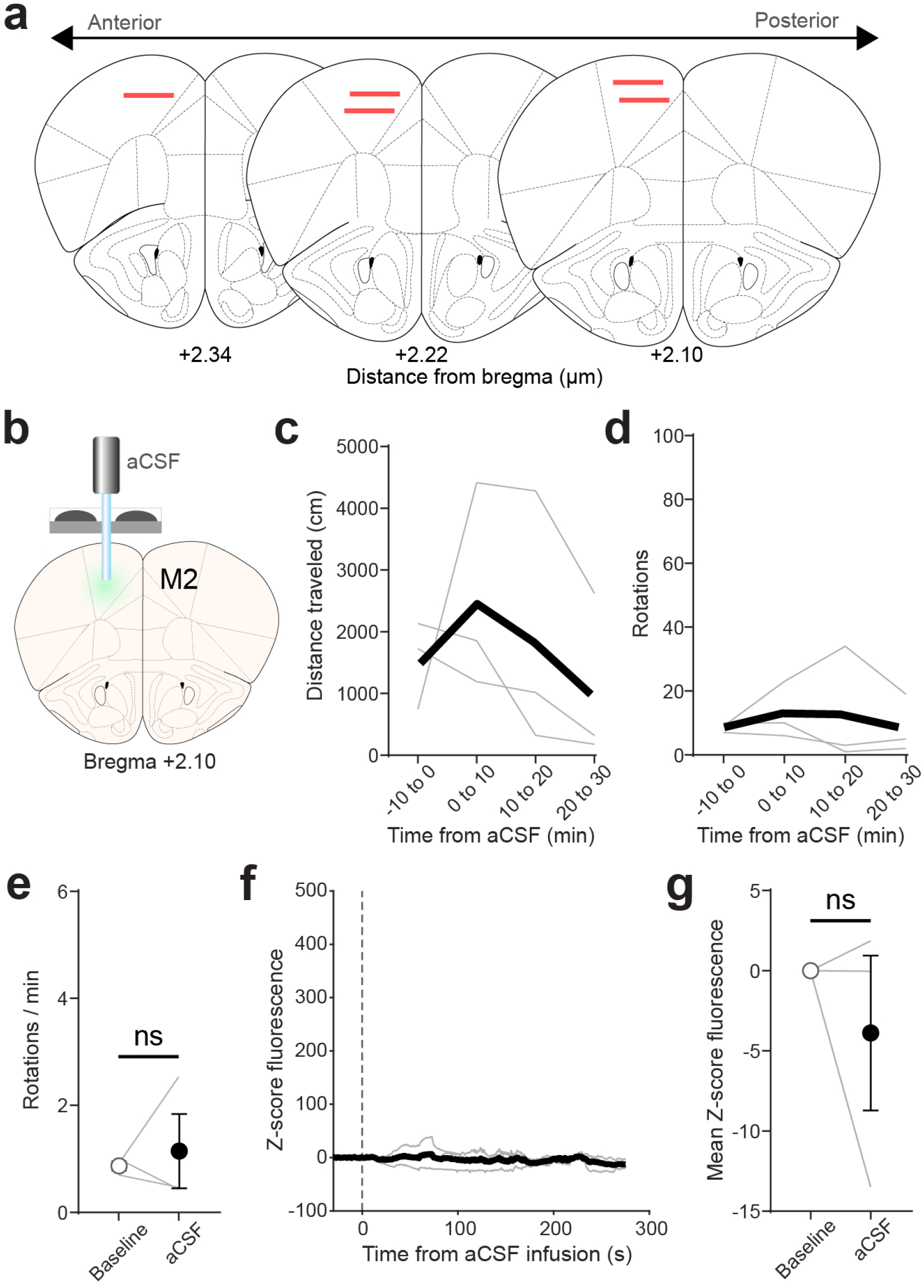
Photo-fluidic infusion of ACSF into M2. (a) Histological verification of photo-fluidic fiber placement (n=5 mice; red line indicates fiber bottom) into M2 at a range of anterior/posterior placements from bregma. (b) Schematic illustration depicting infusion of ACSF into M2. (c) Distance traveled 10 minutes prior to and 30 minutes following ACSF infusion into M2. (d) Number of rotations 10 minutes prior to and 30 minutes following ACSF infusion into M2. (e) No change relative to baseline on the number of rotations per minute following ACSF infusion (paired t-test, t(2)=0.41, p=0.72). (f) No effect of ACSF infusion (dotted line) on GCaMP6 fluorescence into M2. Darker black line indicates mean response across 3 mice. (g) No effect of ACSF infusion on M2 activity as measured by mean GCaMP6 fluorescence (paired t-test, t(2)=0.81, p=0.51).

**Extended Data Figure 7:**
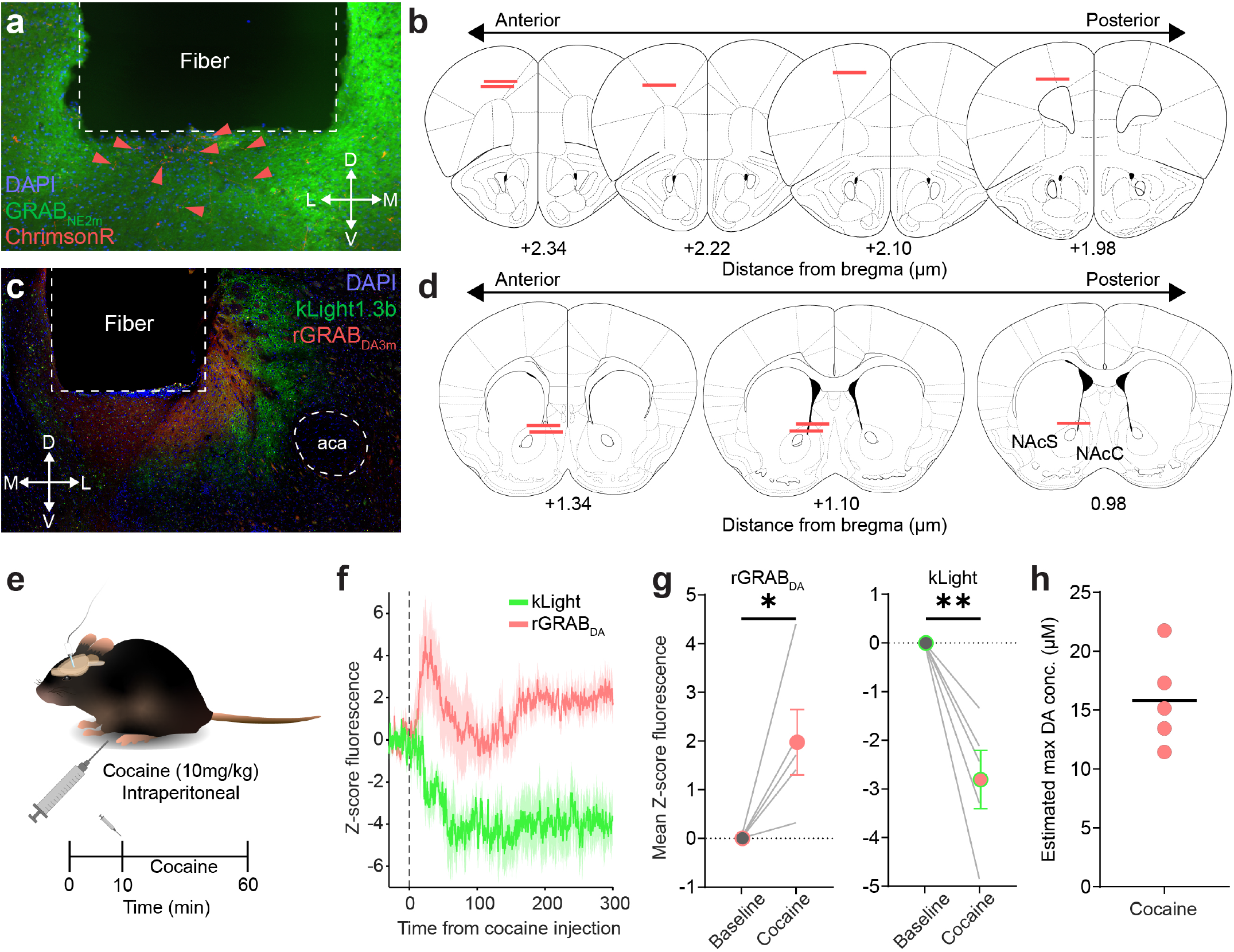
Systemic injection of cocaine while recording NAc GRABDA and kLight signal. (a) High-magnification (20x) image of M2 fiber location above GRAB_NE2m_ (green) and ChrimsonR terminals (red) viral expression. Red arrows indicate ChrimsonR+ terminals. (b) Histological verification of photo-fluidic fiber placement (n=5 mice; red line indicates fiber bottom) into M2 at a range of anterior/posterior placements from bregma. (c) High-magnification (20x) image of NAc fiber location above rrGRAB_DA3m_ (red) and kLight1.3b (green) viral expression. (d) Histological verification of photo-fluidic fiber placement (n=5 mice; red line indicates fiber bottom) into the core of the NAc (NAcC) at a range of anterior/posterior placements from bregma. (e) Schematic illustration depicting intraperitoneal (i.p.) injection of cocaine (10mg/kg) after a 10-minute baseline recording. (f) Z-score fluorescence in response to cocaine injection (n=5 mice). Dotted line indicates i.p. injection of cocaine. Red line represents mean rGRAB_DA3m_ fluorescence (shading indicates SEM) and green line indicates mean kLight fluorescence (shading indicates SEM). (g; left) Systemic injection of cocaine (10mg/kg) increases mean NAc rGRAB_DA3m_ fluorescence relative to baseline (paired t-test, t(4)=2.9, p=0.04). (e; right) Systemic injection of cocaine reduces mean kLight fluorescence relative to baseline (paired t-test, t(4)=4.7, p=0.009). (h) Estimated maximum NAc DA concentration following systemic cocaine injection, based off peak fluorescence obtained following 30µM infusion in Fig. 5d. **p<0.01, *p<0.05. aca=anterior commissure.

